# Conformational dynamics of lipid transfer domains provide a general framework to decode their functional mechanism

**DOI:** 10.1101/2023.04.11.536463

**Authors:** Sriraksha Srinivasan, Andrea Di Luca, Arun T. John Peter, Charlotte Gehin, Museer A. Lone, Thorsten Hornemann, Giovanni D’Angelo, Stefano Vanni

## Abstract

Lipid Transfer Proteins (LTPs) are key players in cellular homeostasis and regulation, as they coordinate the exchange of lipids between different cellular organelles. Despite their importance, our mechanistic understanding of how LTPs function at the molecular level is still in its infancy, mostly due to the large number of existing LTPs and to the low degree of conservation at the sequence and structural level. In this work, we use molecular simulations to characterize dynamical and mechanistic aspects of a representative dataset of Lipid Transport Domains (LTDs) of 12 LTPs that belong to 8 distinct families. We find that LTDs display common dynamical, rather than structural, features despite no sequence homology nor structural conservation. These dynamical features correlate with their mechanistic mode of action, allowing to interpret and design experimental strategies to further dissect their mechanism. Our findings indicate the existence of a conserved, fold-independent mechanism of lipid transfer across LTPs of various families and offer a general framework for understanding their functional mechanism.

## Introduction

Lipids are one of the key building blocks of eukaryotic cells, as they allow for the spatial and temporal organization of chemical reactions in different cellular compartments, called organelles. Eukaryotic cells contain thousands of different lipid types, and each membrane-bound organelle possesses a characteristic lipid composition necessary for its proper functioning^1^. This compositional identity is crucial not only towards their individual functions but also to shape and regulate intracellular signaling and trafficking processes between them^2^.

Since lipid synthesis is not ubiquitous, but rather mostly localized to the endoplasmic reticulum^3^ (ER), lipids must be rapidly transported between organelles to maintain lipid homeostasis and organellar identity. This is achieved via two main routes, the vesicular and non-vesicular pathways. In the vesicular pathway, cargo vesicles, originating from lipid remodeling processes mediated by coat proteins, travel from a donor organelle to an acceptor one, where the vesicle undergoes fusion^4^. This pathway is not only crucial for cellular exocytosis and endocytosis, but also intracellularly along the secretory pathway^4^. Alternatively, in the non-vesicular pathway, trafficking of lipids between organelles is performed by lipid transfer proteins (LTPs), which solubilize lipids and facilitate their transport between two membranes. Non-vesicular lipid transport promotes a more rapid modulation of the lipid composition of organelles compared to vesicular trafficking, and is crucial during stress conditions when vesicular trafficking is compromised^5^.

Despite growing interest in the non-vesicular lipid transport pathway, our mechanistic understanding of how LTPs perform their function is still largely incomplete. The only unifying feature of LTPs is the presence of a lipid transfer domain (LTD) containing a hydrophobic cavity that encloses the lipid, and a polar exterior, making lipid transfer across the cytoplasm a potentially energetically favorable process. Two main models of lipid transport by LTPs have been put forward: the shuttle model and the tunnel model. In the shuttle model, a small (< 50-100 kDa) LTD travels between the donor and the acceptor organelle, cyclically taking up and releasing their substrate lipid^5^. In the tunnel model, a large (>100kDa) LTD physically connects the two organelles, establishing a continuous hydrophobic pathway in which lipids can simply diffuse between the two membranes^5^. In both cases, however, a fine regulation of multiple mechanistic steps must be accurately tuned to achieve lipid transport with the correct directionality and rate, and how such complex coordination is achieved remains largely unclear^5^.

This picture is further complicated by the sheer number of existing LTPs. So far, hundreds of different LTPs, belonging to several distinct protein families^6, 7^, have been identified. This diversity possibly originates from the huge chemical variability of the lipid substrates and organellar membranes they bind to. While this has likely allowed to fine-tune the mechanism and specificity of lipid transport by LTPs, it has so far prevented the establishment of a common framework to understand and interpret the molecular steps underlying this process. To this extent, only few studies have attempted to investigate in a high-throughput fashion functional properties of LTPs, such as their membrane or lipid binding^8^. Rather, the investigation of individual LTPs using cellular biology or reconstitution approaches remains to date the most frequent strategy.

A direct consequence of this case-by-case modus operandi is that a plethora of concurring models have been put forward to explain the mechanism and specificity of lipid transport. Yet, these models are of limited transferability across different LTPs since they largely rely on specific observations on either protein structure (such as the presence of a lid^6, 9^, of electrostatic surface patches^10, 11^ or on the specificity of the lipid-binding cavity^12^) or on experimentally-determined transport properties (such as counter-exchange between different lipid species and lipid-dependent transport rates)^13, 14^.

Here, we employ a computational-based alternative approach to characterize the functional mechanism of LTDs. We find that small LTDs (<50-100kDa) display common dynamical, rather than structural, features. These features correlate with their functional mechanism and suggest a conserved mechanism of lipid transfer across LTPs of diverse families.

## Results

### The membrane binding interface of lipid transport domains does not display any conserved structural signature

To identify common, family-independent properties of LTPs, we opted for an *in silico* analysis based on molecular dynamics (MD) simulations. To this end, we simulated and analyzed 12 LTDs belonging to 8 different families, ensuring a sampling of proteins with diverse secondary structures and partner lipids (Fig. 1a).

**Figure 1:**
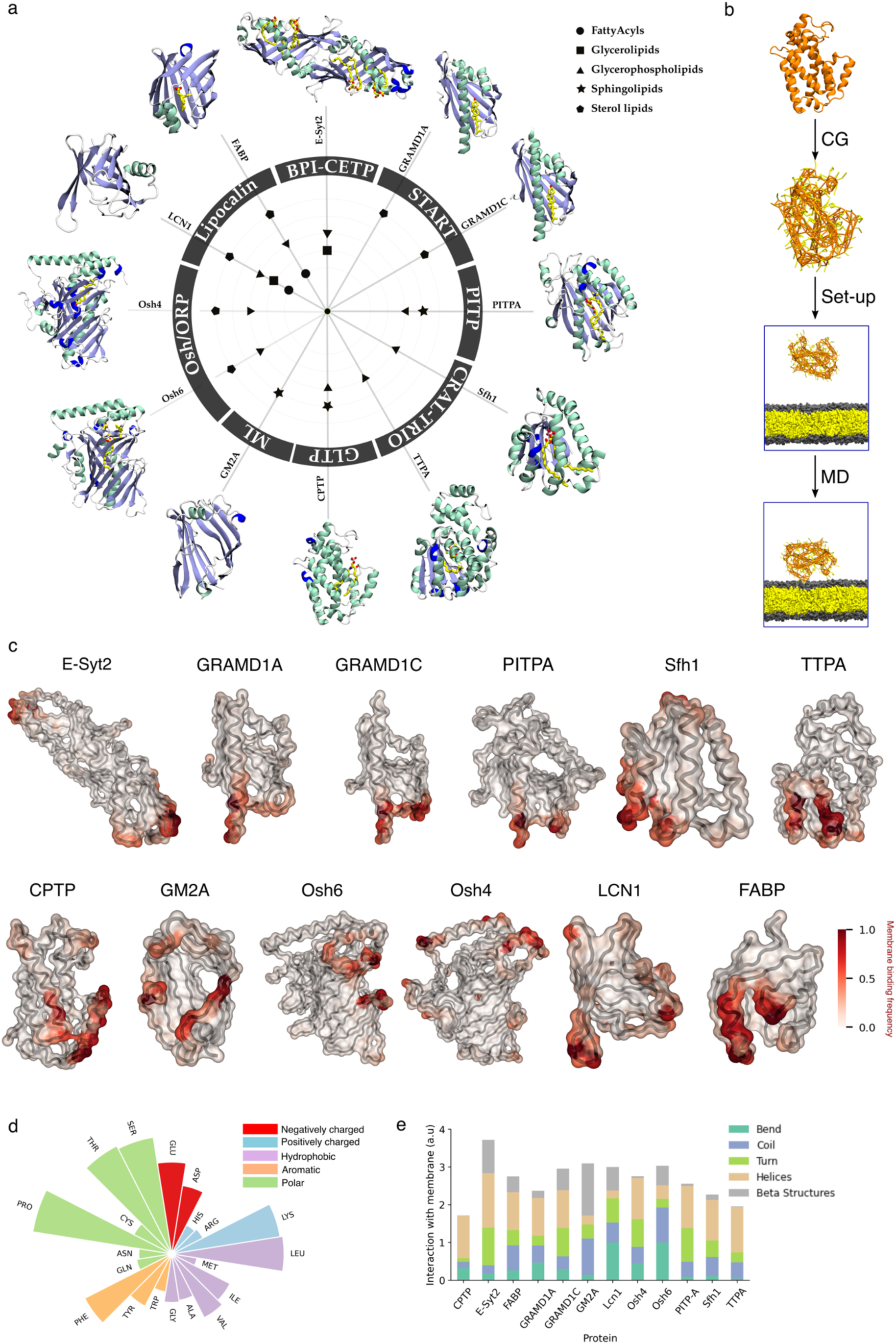
Membrane-binding regions of lipid transfer proteins do not display shared structural features. (**a**) Members of eight different lipid transfer protein families used in this study. Symbols inside the circle indicate their lipid specificity^7^. The secondary structures of the LTDs are shown along the periphery. **(b)** Schematic of the protocol followed to perform MD simulations of LTD-bilayer systems and analyze interacting regions **(c)** LTD structures colored by their frequency of interaction with lipid bilayers **(d)** Interaction frequency of each amino acid with the membrane, summed over all LTDs in the dataset **(e)** Interaction frequency of each secondary structure type. Abbreviations: BPI, bactericidal permeability-increasing protein; CETP, cholesteryl ester transfer protein; CPTP, ceramide-1-phosphate transfer protein; E-Syt2, extended synaptotagmin2; FABP, fatty acid-binding protein; GLTP, glycolipid transfer protein; GM2A, ganglioside GM2 activator protein; GRAMD, Glucosyltransferases, Rab-like GTPase activators and myotubularins domain; LCN1, lipocalin-1; ML, MD-2-related lipid-recognition; ORP, Oxysterol-binding protein (OSBP)-related protein; PITPA, PI-transfer protein; START, StAR-related lipid-transfer; TTPA, tocopherol transfer protein-A.

Since LTDs uptake lipids from membranes of cellular organelles, we first investigated how different LTDs bind to model lipid bilayers, using a computational protocol we recently developed^15^. In short, to determine the membrane-binding interfaces of LTDs, we performed coarse grain (CG) simulations of the proteins in combination with pure dioleoyl-phosphatidylcholine (DOPC) lipid bilayers, starting with the protein at a minimum distance of 3 nm away from the bilayer (Fig. 1b). Over the course of the MD simulation, the proteins exhibit transient interactions, *i.e.* multiple binding and unbinding events, as shown by the minimum distance curves in Fig. S1. The membrane-interacting residues of each protein are then determined by computing the frequency of the interaction of each residue with the bilayer (see Methods).

Fig. 1c shows the structure of each protein in our dataset, colored by the frequency of interaction with the lipid bilayer. The corresponding residue-wise analysis of the frequency of interactions is reported in Fig. S2. The excellent agreement between the membrane interface determined from the simulations and the experimentally-proposed one available for CPTP^16^, FABP^17^, Osh4^18^, Osh6, PITPA^19^, and TTPA^20^ (Fig. S2) suggests that our *in silico* methodology is able to reproduce the correct membrane binding interface of LTDs, similar to what was shown for different families of membrane-binding peripheral proteins^15^.

Using the obtained residue binding propensity, we next investigated whether LTDs belonging to different families possess common structural membrane binding signatures. First, we investigated whether the membrane binding interface displays specific sequence properties (Fig. 1d and S3). Concomitant analysis of all LTDs (Fig. 1d) indicates that the membrane binding interface of LTDs is enriched in positively charged amino acids (Lys) and aromatic/hydrophobic ones (Phe, Leu, Val, Ile). This confirms previous observations, as (i) binding of negatively charged lipids via positively charged residues and (ii) hydrophobic insertions are two of the main mechanisms involved in membrane binding by peripheral proteins^21–25^. Most notably, also polar residues such as Ser, Thr, and Pro seem to be involved in membrane binding by LTDs. However, analysis on a protein-by-protein scale reveals a lack of any general trend, as each protein family displays a characteristic amino acid composition in its membrane binding interface (Fig. S3). Similarly, analysis of the secondary structure of the membrane binding interface does not display any preference for secondary structure elements (Fig. 1e).

#### The membrane binding interface of lipid transport domains is characterized by large collective motions

We next investigated whether dynamical properties, rather than structural ones, could explain, or correlate with, membrane binding in LTDs. To do so, we performed extensive atomistic MD simulations of the proteins in solution, and we characterized their dynamical signature using principal component analysis (PCA) (Fig. 2a). In this technique, thermal and high-frequency fluctuations occurring during protein dynamics are filtered out to identify the dominant large-scale and slower motions of the protein^26, 27^. Using this approach, we could identify the protein regions undergoing the largest collective motion (Fig. S4).

**Figure 2:**
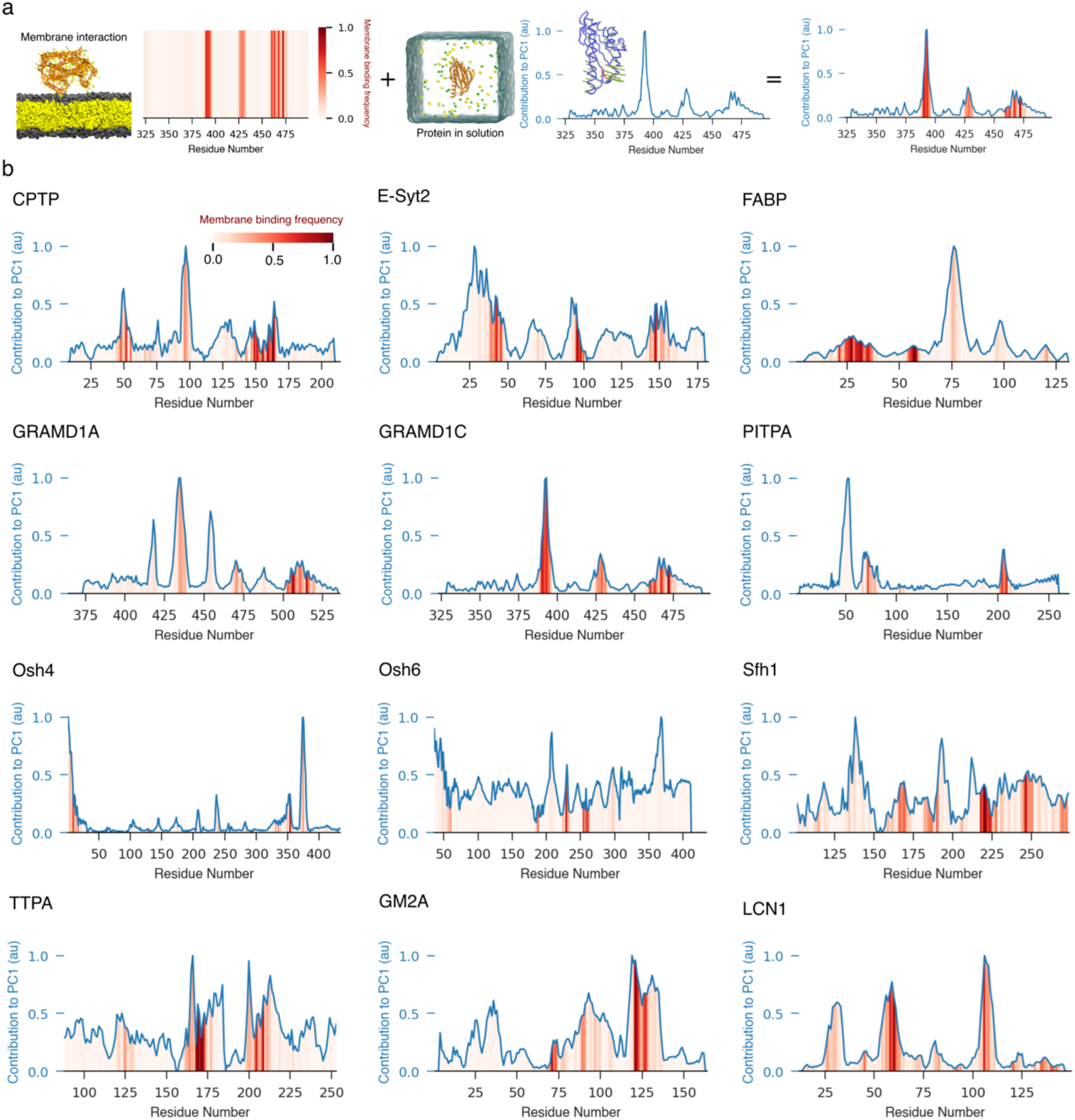
Membrane-interacting regions of LTDs correlate with their dynamical regions. **(a)** Schematic of the combination of membrane-binding interface of the protein from CG-MD simulations and the dynamics of the protein in water from atomistic MD simulations **(b)** The frequency of interaction of each residue with the lipid bilayer is represented by the heatmap in red while the contribution of each residue to the first principal component of protein dynamics (PC1) is indicated by the line plot in blue. Both measures have been normalized to a value between 0 and 1 for each protein, respectively.

We next combined this analysis with our membrane-binding assay (Fig. 2a), to investigate whether membrane-binding regions are also involved in the proteins’ principal motions. Notably, we observed that almost all LTD membrane binding regions (red bars, Fig. 2b) correspond to maxima of the contributions of single-residue dynamics to PC1 (blue curves, Fig. 2b). This indicates that protein regions located at or near the membrane-binding interface display large collective motions regardless of the overall protein fold and structure. While not all PC1 maxima are membrane-binding regions, which is to be expected since PCA describes collective motions that could be at distant protein locations, our data nevertheless suggest that the dynamical signature of LTDs provides information on their membrane binding properties.

#### Lipid transport domains in solution display a conformational equilibrium that is modulated by the presence of bound lipids

Since our data indicate a correlation between LTDs’ collective dynamical properties and membrane binding, we next opted to further characterize the dynamical conformational landscape of LTDs. To do so, we computed the population distribution of the projections of the first principal component (PC1) for all LTDs in our dataset, and we clustered the resulting conformations using a density-based automatic procedure^28^ (Fig. 3a,b). Our results indicate that in the apo form, the proteins sample a diverse conformational landscape as shown by the multimodal distribution of the populations of PC1 (Fig. 3c, orange histograms). Notably, and despite significant differences in the conformational landscape of the various LTDs, density-based automated clustering is able to distinguish at least two distinct clusters for each protein (Fig. 3c, bar plots).

**Figure 3:**
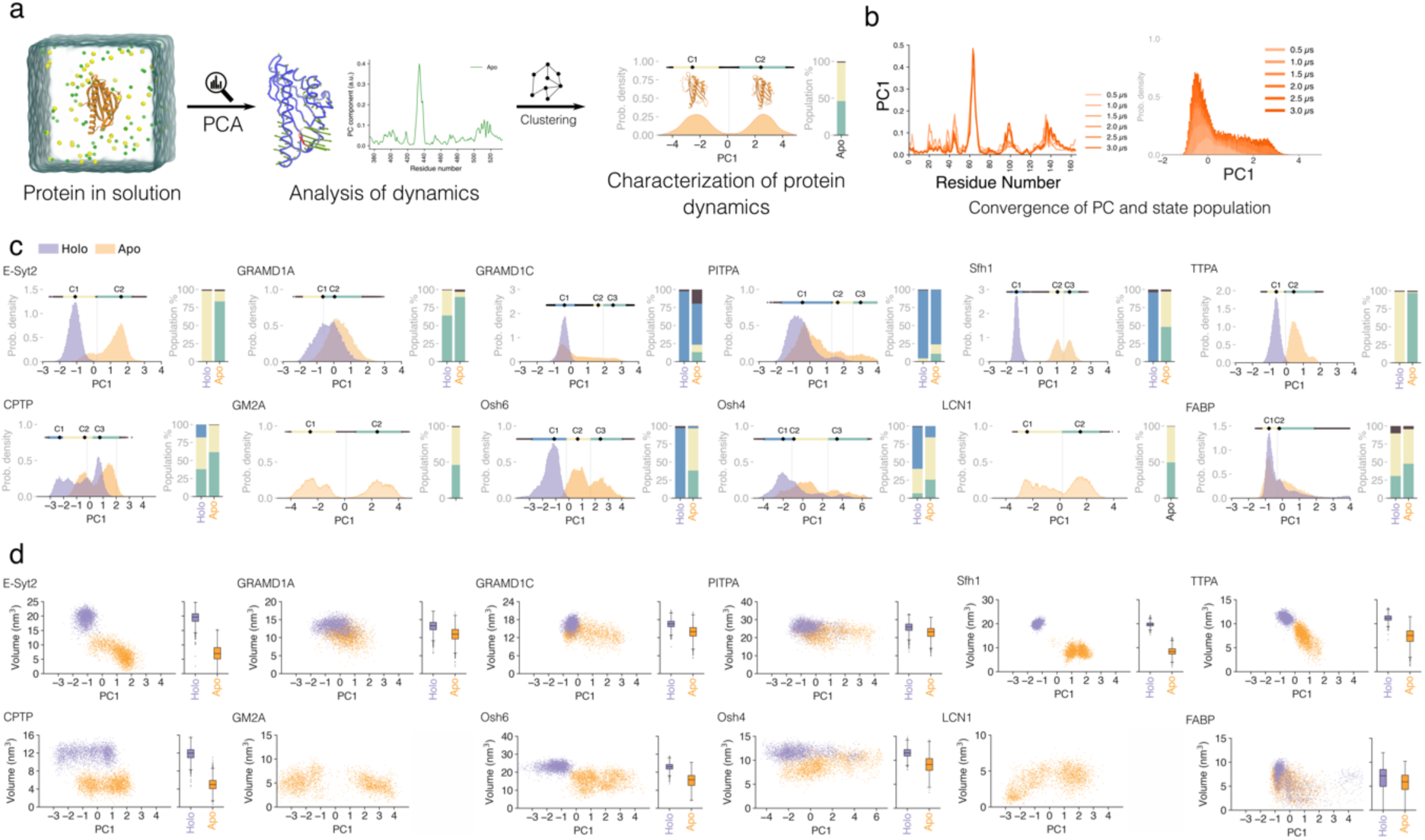
Bound lipids modulate the conformational landscape of LTDs. **(a)** Protocol to characterize protein dynamics from atomistic simulations of the protein in solution. The first principal component PC1 was determined, and the resulting conformations were clustered using a density-based automatic procedure. **(b)** Simulations of 6 replicas for 500 ns each were found to be sufficient to ensure convergence of the distribution of PC1. **(c)** The population distributions of PC1 from apo (orange) and holo (purple) simulations of the protein are shown. The clusters of the distributions are indicated by the line above the histogram, with the black dot (labelled C1, C2, C3) representing the cluster center. Bar plots on the right indicate the relative apo and holo populations of each cluster. Holo-forms of the protein could not be simulated for GM2A and LCN1 due to the lack of lipid-bound crystal structures. **(d)** Cavity volumes for apo (orange) and holo (purple) forms of the protein, computed on the projections of PC1.

The dynamical alternation between different conformations is a hallmark of enzyme dynamics, where it generally correlates with protein activity^29^. To push further this parallelism, we next investigated the effect of the bound lipid on LTD dynamical behavior, since in classical enzymology the presence of a bound substrate is generally known to restrict enzyme dynamics and stabilize the protein in a specific conformation^30^.

To determine if the presence of the bound lipid would alter the dynamics of the LTDs in our dataset, for all cases in which a bound lipid was co-crystallized together with the protein (10 out of 12 proteins in our dataset), atomistic simulations in the holo-form were performed and analyzed following the same protocol used for the apo-forms. Notably, the comparison of the projections along the PC1 from the holo-form with that of the apo-form shows that, for all proteins, the presence of the bound lipid shifts the population distributions along the PC (Fig. 3c, purple histograms). The residue-wise contribution to PC1 for simulations in the apo form, holo form, and when taken together is shown in Fig. S4. In detail, the presence of a bound lipid appears to stabilize the protein in one specific conformation (Fig. 3c, bar plots). In most cases, this conformation is also sampled by the protein in its apo form, but always with a lower frequency than the corresponding holo simulations, indicating that the presence of a bound lipid indeed “locks” the LTD in a specific 3D structure, akin to substrate binding in enzyme dynamics.

To further characterize this conformational landscape, we next evaluated the structural properties of representative conformers belonging to the two main clusters emerging from the PC analysis. To do so, we first computed the root mean square deviation (RMSD) of structures corresponding to the two extremes of the clusters along the PC1 spectrum (Fig. S5). For the LTDs in our dataset, this value varies between 2.3 and 7.1 Å, indicating that the range of structural difference between clusters is highly variable.

Next, we computed the volume of the hydrophobic cavity of the structures sampled in our MD simulations for both apo and holo trajectories. For almost all LTDs investigated here, we could quantify significant differences in cavity volume for the two conditions (apo *vs* holo) (Fig. 3d). In almost all cases, when no lipid is found inside the protein, its hydrophobic cavity shrinks, thus reducing its size (Fig. 3d). Consistent with the behavior of the entire protein (Fig. 3c), the cavity in the holo state generally adopts a more compact distribution of conformations (Fig. 3d, purple), while, in the absence of bound lipids, it is able to sample multiple conformations (Fig. 3d, orange), often including those that are representative of the holo state.

#### The conformational equilibrium of lipid transport domains modulates their lipid transport activity in cells

Our data suggest that all LTDs exist in an equilibrium between two or more conformations, and that the presence of bound lipids alters their equilibrium population, akin to a mechanism of “conformational selection” or “induced fit”. For all LTDs, these conformational changes are localized in protein regions that interact with the lipid bilayer. Since the main proposed activity of LTDs is to transport lipids, we next sought to investigate whether this observed “enzyme-like” behavior has functional consequences.

We first sought to investigate whether the observed conformational dynamics of LTDs can provide clues into experimentally determined lipid transport rates. To this extent, the two members of the ASTER family in our dataset provide a fertile ground for this analysis, since the ASTER domains of GRAMD1A and GRAMD1C have very similar structures (RMSD X-ray=0.76Å) yet very different transport rates^31^, with GRAMD1A transporting ∼8.4 and GRAMD1C between ∼0.5 and 1.5 DHE molecules/min/molecule LTP in identical experimental conditions *in vitro* (Fig. S6)^31^. Notably, despite their structural similarities, the two proteins display markedly different conformational populations in our dynamical analysis, with GRAMD1A exhibiting a significantly more overlapped population between apo and holo states compared to GRAMD1C (Fig. S6). This observation correlates well with the ability of GRAMD1A, but not of GRAMD1C, to interchange between different conformations, suggesting that overlap between apo and holo states could promote lipid transport activity.

Next, we investigated whether mutations able to alter the conformational dynamics of LTDs, as suggested by our MD simulations, would also impair lipid transport *in cellulo*. To do so, we selected two proteins that were not in our original dataset, but for which lipid transport activity has been extensively characterized in cellular assays: STARD11 (also known as CERT), a member of the START family^32^, and Mdm12, a member of the BPI-CETP/SMP/TULIP family and a part of the ERMES complex^33, 34^.

STARD11 is an LTP that promotes ceramide transport from the ER to the *trans*-Golgi at ER-Golgi membrane contact sites. It is composed of an N-terminal Pleckstrin Homology (PH) domain mediating its anchoring to the *trans*-Golgi^35, 36^, a two phenylalanines in an acidic tract (FFAT) motif responsible for binding to the ER membranes^35^ and a C-terminal and a (StAR)-related lipid transfer (START) domain that extracts ceramide from the ER membrane and delivers it to the *trans* Golgi. Once transported by STARD11 to the *trans*-Golgi, ceramide is readily converted into sphingomyelin^37^.

To alter the conformational dynamics of the START domain of STARD11, we mutated the Ω1 loop (Fig. 4a), that is involved in the principal motion of the protein for both STARD11 (Fig. S7a) as well as for the other START-containing LTDs in our dataset, GRAMD1A and GRAMD1C (Fig. 2). Analogously to what is proposed for ASTER proteins^38^, we opted to mutate residues 497-501 (RVWPA, which are residues 471-475 in the crystal structure) into a stretch of prolines (for short, STARD11-5P) (Fig. 4a). As expected, this mutation results in a significant shift of the population distribution of STARD11 along the first principal component (Fig. S7b).

**Figure 4:**
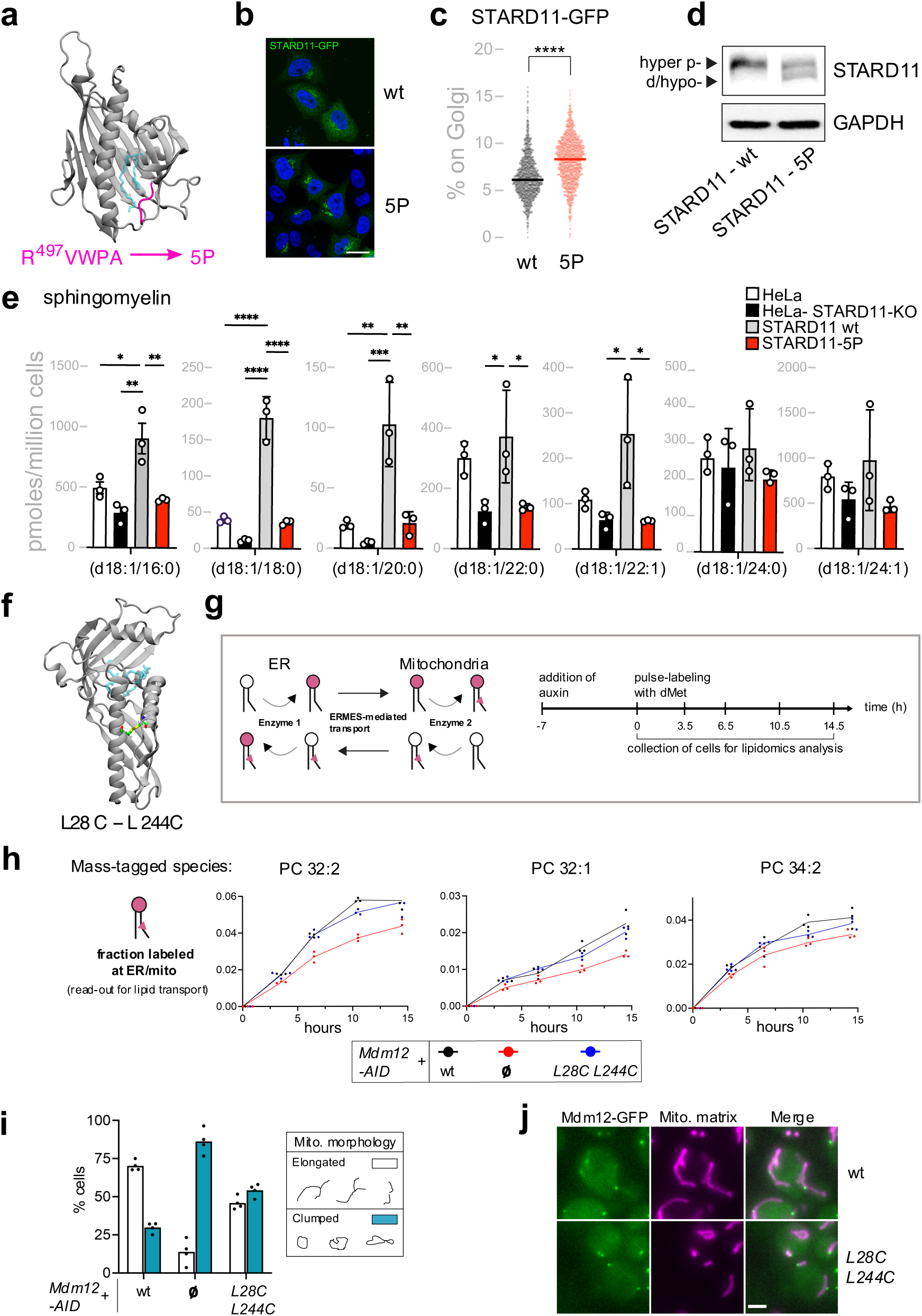
Mutations altering LTD dynamics impair their lipid transport properties. **(a)** Design of the STARD11-5P mutant at the omega loop (pink). The position of the lipid in the crystal structure is shown in transparent blue licorice. **(b)** STARD11-GFP WT (wt) and the STARD11-5P-GFP (5P) mutant localization in HeLa cells analysed by confocal microscopy. Scale bar, 20 μm. **(c)** Percentage of STARD11-GFP WT (wt) and the STARD11-5P-GFP (5P) mutant associated with the Golgi complex in HeLa cells. Cells were stained with Hoechst and anti-GM130 antibody and analysed by automated fluorescence microscopy (n>1,000 cells per condition; ****p <0.0001 [Students-T-Test]. STARD11-GFP WT grey, STARD11-5P-GFP in red). **(d)** Western blot of HeLa cells expressing STARD11-GFP WT (wt) or the STARD11-5P-GFP (5P) mutant. Hyperphosphorylated (hyper-p-) and de/hypophosphorylated (d/hypo-) bands are indicated by arrowheads. **(e)** Mass spectrometry profile of major sphingomyelins in HeLa cells; Hela cells *STARD11*-KO, and Hela cells *STARD11*-KO overexpressing STARD11-GFP WT (wt) or the STARD11-5P-GFP (5P) mutant. (n=3; data are means ± SD; *p <0.05, **p <0.01; ***p <0.001 [Ordinary one-way ANOVA]) **(f)** Design of the Mdm12 L28C-L244C disulfide bridge mutant with sulfur shown in yellow, carbon in green, oxygen in red, and hydrogen in white. The position of the lipid in the crystal structure is shown in transparent blue licorice. **(g)** METALIC uses the endogenous methyltransferases Cho2/Opi3 (Enzyme 1) localized to the ER to mass tag the headgroup, and the mitochondria matrix-targeted CFAse (Enzyme 2), a bacterial enzyme, to mass tag the fatty acid chains in phospholipids. Doubly mass tagged lipids (pink head group and fatty acid tail) serve as a read-out for ER-mitochondria lipid exchange (left panel). Scheme showing time points of auxin and dMet (deuterated methionine) addition. Deuterated *S*-adenosyl methionine resulting from dMet serves as the cofactor for both enzymes (right panel). **(h)** Line plots showing the doubly labeled fraction over time in the Mdm12-AID strain bearing the indicated plasmids. Ø denotes empty vector. Three independent clones of each genotype were used. **(i)** Bar plot depicting percentage of cells with the classified mitochondria morphology in the Mdm12-AID strain – after 5 h auxin treatment – bearing the indicated plasmids. N = ∼100 cells for each condition from four randomly chosen regions of interest. **(j)** Localization of C-terminally GFP-tagged Mdm12. Mitochondria-matrix targeted CFAse-mCherry is used as a mitochondria marker. Scale bar, 2 µm.

Next, to test STARD11-5P bioactivity we first expressed GFP-tagged STARD11-5P and compared its intracellular localization with that of STARD11 wildtype (wt). Intriguingly, we found that STARD11-5P is significantly more membrane associated than STARD11 wt (Fig. 4b, c). STARD11 membrane association is controlled by its phosphorylation status with hyper and hypo phosphorylated forms of STARD11 being more cytosolic and membrane associated, respectively^39, 40^. Accordingly, we found that STARD11-5P is hypo-phosphorylated relative to STARD11 wt (Fig. 4d).

Modulation of STARD11 phosphorylation/ localization is part of a homeostatic circuit whereby excessive STARD11 activity and sphingomyelin production trigger a signaling reaction leading to its phosphorylation and inactivation^39, 41, 42^. STARD11-5P appears to be unable to induce this response possibly due to its inability to sustain ceramide transfer and conversion to sphingomyelin. To test this hypothesis, we evaluated the sphingomyelin levels in STARD11-KO HeLa cells^37, 39^ expressing either STARD11 wt or STARD11-5P. We found that while expression of STARD11 wt rescues the defect in sphingomyelin production in STARD11-KO HeLa cells, STARD11-5P fails to do so (Fig. 4e). Thus, impairing the dynamics of the START domain of STARD11 inhibits its bioactivity.

We next applied a similar approach for Mdm12, a member of the BPI-CETP/SMP/TULIP family of lipid transporters. We wondered if impairing its transition to the “holo-like” conformation would result in defective lipid transport. To do so, we identified protein regions in Mdm12 that are involved in the transition from “apo-like” to “holo-like” based on the analysis of the PC1 motion of E-Syt2, the other protein with a BPI-CETP/SMP/TULIP domain in our dataset (Fig. 3). The dominant motion in the SMP domain of E-Syt2 consists of a significant breathing motion of the hydrophobic cavity, with the cavity doubling its size in the presence of a bound lipid (Fig. 3d). By structural analogy between E-Syt2 and Mdm12, we identified a pair of residues (L28, L244) in Mdm12 that could undergo significant variations in their relative distance amongst the “apo-like” and the “holo-like” conformations. We thus decided to mutate them into cysteine residues to promote the formation of a disulfide bridge that is expected to lock the protein in the “apo-like” state (Fig. 4f). This mutation also results in a shift in the conformational population along PC1 for Mdm12 (Fig. S7c, d).

To directly quantify lipid transport activity by Mdm12 as part of the ERMES complex, we took advantage of METALIC, a recently developed mass spectrometry based approach that uses enzyme-mediated mass tagging of lipids at two chosen organelles as a proxy to assess lipid transport *in vivo*^43^ (Fig. 4g). Using this method, it was demonstrated that the ERMES complex plays a major role in the transport of phospholipids between the ER and mitochondria^43^. Notably, this protein complex contains two other LTDs, Mmm1 and Mdm34, apart from Mdm12 that possibly arrange in a sequential fashion to achieve lipid transport between the two organelles^33^. Loss of any one of the ERMES subunits renders the whole complex non-functional^34^. To assess the role of the Mdm12 (*L28C L244C*) mutant in ER-mitochondria lipid exchange, we expressed it in a strain where the endogenous Mdm12, fused to auxin-inducible degron (AID), can be inactivated in the presence of auxin. In the *L28C L244C* mutant, the lipid transport kinetics was similar to the wild-type (wt) for the assessed phospholipid species except at t=10.5h where there was a mild but significant reduction in the double mass labeling of PC 32:2 (12%, *P*=0.002), PC 32:1 (17%, *P*=0.04) and PC34:2 (16%, *P*=0.031) (Fig. 4h). This rather modest phenotype in lipid transport however seemed to manifest its effect on mitochondria morphology. ERMES null mutants bear a clumped mitochondria phenotype rarely seen in wt cells, which mostly have an elongated morphology^44^. Strikingly, the *L28C L244C* mutant manifested an intermediate phenotype with ∼50% of cells bearing clumped mitochondria (Fig. 4i,j). To confirm that the phenotype was not due to alternative reasons, we verified that the *L28C L244C* mutant was expressed, and that it localized similar to the wt version as foci proximal to mitochondria (Fig. 4j).

Taken together, these observations suggest that the conformational alterations observed in our MD simulations upon mutations in STARD11 and Mdm12 are physiologically relevant, and further support our model that the dynamic structural transitions of LTPs are important for their activity and cellular function.

## Discussion

Based on experimental observations on individual LTPs^10, 19, 45–49^, several concurring models have been put forth in recent years to elucidate their mechanism of action^6, 50, 51^. Even though these models have highlighted active-site residues and membrane binding regions or protein sequences that act as potential gates for lipid uptake/release, a unifying model to explain LTP function in a global context remain lacking. Building on this limitation, in this work, we used MD simulation to investigate several LTPs concurrently to decipher the common thread in their mechanism of action. First, using our previously established protocol to study peripheral protein-membrane interactions, we determined the membrane binding interface of 12 LTDs that belong to different families and have diverse secondary structures. We observed a lack of commonality at their membrane binding interface in terms of structural properties such as the amino acid and secondary structure composition. However, we found that across LTDs, the interface and/or residues adjacent to it exhibited high dynamics in solution, and that they displayed the largest collective motion. This observation hints that, irrespective of the nature of the transported lipid, the high degree of protein dynamics we observed in membrane proximal regions could potentially facilitate the energetically unfavorable reaction that involves the extraction of a lipid from a membrane and its transfer to the hydrophobic groove of a LTP. Notably, lipid desorption from the membrane has been proposed as the rate-limiting step for sterol transport proteins^52^.

Furthermore, we established that the dynamics of LTDs can be modulated by the bound lipid, and this modulation alters the conformational preference of the LTD. Our results generalize previous observations made by us^33^ and others^31, 53–56^ in the context of individual LTPs. We foresee that future studies will focus on the functional consequence of such observation, and most notably to the characterization of the extent to which such conformational changes affect multiple steps of protein function, including membrane binding or lipid extraction and release.

Mutations that alter lipid transport have often been proposed to do so by changing the affinity for the membrane due to a change in the membrane binding interface^10, 57, 58^. A key hypothesis arising from our findings is that mutations can also abolish or decrease lipid transfer by affecting the ability of the protein to transition between apo-like and holo-like conformations. Of note, this mechanistic hypothesis is consistent with the one proposed for several Osh/ORP proteins in which experimental deletion of the N-terminal lid results in defective membrane association and/or lipid transport^10, 54, 59, 60^. Moreover, we observe that several mutations that have been shown to alter lipid transfer for some LTDs in our dataset such as Osh6 L69D^10^, Osh4 τι69^54^, PITPA P78L^61^, GRAMD1A 01-5P^38^, and CPTP V158N^11^ are all located in protein regions that our simulations suggest being involved in protein dynamics and membrane binding (Fig. 2b, Fig. S2).

Despite providing structural and dynamical insights into the mechanism of LTPs, our approach has limitations. First, our atomistic sampling of the conformational dynamics of LTDs is finite, and even though PCA analysis allows to describe the essential dynamics of the LTDs, we can’t completely describe larger conformational changes that might occur on longer timescales, such as, for example the opening/closing of the lid occurring in Osh proteins upon membrane binding^10, 53, 54^. Second, our membrane binding assay, based on CG simulations with the MARTINI model, has two major limitations: it requires the use of an elastic network to restrain the secondary structure of the protein^62^ and it lacks sensitivity towards the lipid composition of membranes. Hence, this method is not well suited to investigate whether the conformational changes induced by the presence of the lipid inside the LTD cavity alter the membrane binding propensity and to what extent membrane properties such as electrostatics, curvature, and packing defects play a role in the directionality of the transfer cycle. Deciphering the lipid transfer reaction *in silico* remains a challenging task, as several mechanisms including protein dynamics, protein-membrane interactions and protein-protein interactions contribute to the overall process. However, our approach to understand LTD dynamics has greatly facilitated the identification of physiologically relevant regions/amino acids in LTPs that underlie their activity (Figure 4).

Overall, our work demonstrates the importance of conformational dynamics of LTPs in their mechanistic mode of action and provides a framework to understand their functional mechanism more generally, in a protein-aspecific context. The observation that all LTDs share dynamical features raises interesting questions concerning the evolutionary origin of this apparent convergence. We hope our results will contribute to the progress of this still underappreciated aspect of LTP research.

## Methods

### Coarse-Grain Simulations of protein-membrane systems

The atomistic structure of the LTDs studied were obtained from RCSB PDB^63^ (Table 1) and were converted to a CG model using the martinize script. An additional elastic network with a force constant of 700 kJ mol^-1^ nm^-2^, and upper elastic bond cut-off of 0.8 nm, was used to restrain the secondary structure of the protein. The CHARMM-GUI Bilayer Builder for Martini^64^ was used to build lipid bilayers with lateral dimensions of 20 nm x 20 nm. The bilayers were then equilibrated according to the standard six-step equilibration protocol provided by CHARMM-GUI. Water molecules and ions were stripped off the system, and the protein of interest was placed away from the membrane, such that the initial minimum distance between any CG bead of the protein and any CG bead of the membrane was at least 3 nm. The orientation of the protein was such that its principal axes were aligned with the *x,y,* and *z* directions of the system, with the longer dimension of the protein along the *z* direction. This setup was then solvated and ionized with 0.12 M of sodium and chloride ions to neutralize the system and reproduce a physiological salt concentration.

**Table 1:** LTDs used for MD simulations.

| Protein | PDB ID | Residues of LTD used (Uniprot numbering) | Bound-lipid in atomistic simulations of holo-form |
| --- | --- | --- | --- |
| CPTP | 4KBS (apo),<br>4K8N (holo) | 8-214<br>8-214 | Ceramide-1-phosphate |
| E-Syt2 | 4P42 | 191-370<sup>a</sup><br>(chain A and B) | Dioleoylglycerol<br>(2 molecules per chain) |
| FABP | 5CE4 | 2-132 | Oleic acid |
| GRAMD1A | 6GQF | 364-536 | Cholesterol |
| GRAMD1C | 6GN5 | 323-499 | Cholesterol |
| Osh4<sup>d</sup> | 1ZHZ | 1-434 | Ergosterol |
| Osh6 | 4B2Z | 36-434 | DOPS |
| PITPA | 1T27 | 2-270 | DOPC |
| Sfh1 | 3B74 | 101-274<sup>b</sup> | DOPE |
| TTPA | 1OIP<sup>**</sup> | 88-253<sup>b</sup> | α-tocopherol |
| GM2A | 1PU5 | 33-193 | - |
| LCN1 | 1XKI | 30-168 | - |
| STARD11 | 2E3R | 364-624 | - |
| Mdm12 | 5GYD | 1-268<sup>c</sup> | - |

DOPC: Dioleoyl-phosphatidylcholine; DOPE: Dioleoyl-phosphatidylethanolamine; DOPS: Dioleoyl-phosphatidylserine
<sup>a</sup>Only one of the two chains was considered for analysis of the atomistic simulations.
<sup>b</sup>Additional residues (Sfh1: 4-310 and TTPA: 25-275) were retained in the simulations in order to maintain the secondary structure of the protein, but were not considered for the analysis since they do not constitute the CRAL-TRIO lipid transfer domain.
<sup>c</sup>The loop missing from the crystal structures deposited in the RCSB PDB structure 5GYD was rebuilt by adding 5 residues to each truncated side of the structure. The loop has been shown to be dispensable for function<sup>71</sup>.
<sup>d</sup>The PDB structure of Osh4 in complex with cholesterol (1ZHY) was used for coarse-grained simulations with the bilayer.

8 independent replicas of 3 μs each were simulated for each LTD-membrane system using the GROMACS^65^ (2018.x-2021.x) package and the Martini 3 force field^66^. Energy minimization was performed using the steepest descent algorithm, followed by a short MD run of 250 ps. Production runs were performed at a temperature of 310K using a velocity-rescale thermostat^67^, with separate temperature coupling for protein, bilayer, and solvent particles and a coupling time constant of 1.0 ps. The md integrator was used for the production runs, with a time step of 20 fs. The Parrinello-Rahman barostat ^68^ was used to maintain the pressure at 1 bar, with a semi-isotropic pressure coupling scheme and a coupling time constant of 12.0 ps. The Coulombic terms were computed using reaction-field ^69^ and a cut-off distance of 1.1 nm. A cutoff scheme was used for the vdW terms, with a cut-off distance of 1.1 nm and Verlet cut-off scheme for the potential-shift^70^. The Verlet neighbor search algorithm was used to update the neighbor list every 20 steps with a buffer tolerance of 0.005 kJ mol^−1^ ps^−1^. Periodic boundary conditions were used in all three directions. The system setup and simulation parameters are in accordance with the recently proposed protocol to study transient protein-membrane interactions with the Martini force field^15^.

### Atomistic Simulations of protein in solution

The CHARMM-GUI Solution Builder^72^ was used to construct set ups of the protein in solution in the apo (without lipid) and holo (lipid-bound) forms as indicated in Table 1. The stoichiometry of protein to lipids were one-to-one for all LTDs in the dataset except for the SMP domain of E-Syt2. The electron density maps of the SMP domain show clear densities for two lipid-like molecules per E-Syt2 monomer^73^. One density is consistent with that of a dioleoylglycerol lipid (DOG), while the other is reminiscent of Triton X-100, a detergent used in protein purification. Denaturing mass-spectrometry however shows that two lipid molecules can bind per monomer^73^. Hence, we replaced the two detergent molecules in the domain with a DOG each, by aligning a vector from the head-group to tail of the reminiscent detergent with that of a DOG molecule. Thus, in total, 4 molecules of DOG were used for simulations of the E-Syt2 SMP in the holo-form. Standard CHARMM-GUI lipid parameters were used for DOG, oleic acid, cholesterol, ergosterol, DOPC, DOPE, and DOPS lipids, while parameters for ceramide-1-phosphate were obtained from the CGenFF web server^74^ and parameters for α−tocopherol were determined by using the structure density factors in the PDB structure when constructing the system using the CHARMM-GUI Solution Builder. A cubic box of edge length was constructed such that the distance between the protein and the edge of the box was at least 1 nm, and the systems were solvated with TIP3P water and ionized with 0.12M of sodium and chloride ions. 6 independent replicas of 500 ns each were simulated for each system using the GROMACS^65^ (2018.x-2021.x) package and the CHARMM36m force field^75^. Energy minimization was performed using the steepest descent algorithm and was followed by a short NVT and NPT equilibration of 100 ps each with position restraints on the backbone atoms of the protein. Production runs were performed using a velocity-rescale thermostat^67^ at a temperature of 310K, with separate temperature coupling for protein and solvent particles, and a coupling time constant of 0.1 ps. The first 10 ns of the production runs of each replica were not considered for analysis. The md integrator was used for the production runs, with a time step of 2 fs. The pressure was maintained at 1 bar by using the Parrinello-Rahman barostat^68^ and an isotropic pressure coupling scheme with a compressibility of 4.5 x 10^-5^ bar^-1^ and a coupling time constant of 2.0 ps. The Particle Mesh Ewald (PME) methods was used to compute the electrostatic interactions with a Fourier spacing of 0.16nm, a cutoff of 1.2 nm, and an interpolation order of 4 was used. Van der Waals (VDW) interactions were switched to zero over 10 to 12 Å. The LINCS algorithm was used to constrain bonds involving hydrogen atoms. Periodic boundary conditions were employed in all three directions.

To confirm the conformational differences between apo and holo states, simulations of the protein in the apo-form were performed for E-Syt2, Sfh1, and TTPA, by starting from the final structure obtained from the holo simulations by removing the lipid from the last frame of the simulations (500 ns x 6 replicas). The sampling and PC obtained in this case are similar to the ones obtained from simulating the apo form of the LTD starting from the crystal structure, indicating that all conformational changes observed are reversible and the protein is not trapped in a metastable state (Fig. S8).

### Mutant simulations

#### Mdm12

Predictions for the disulfide bridges mutations were based on the Mdm12 structure by taking pairs of residues with alpha-carbon atom distances below 8 Å. The mutations to cysteine and the construction of the disulfide bridge were done using the CHARMM-GUI Solution Builder.

*STARD11.* The 5P mutation of the START domain of STARD11 was done using the CHARMM-GUI Solution Builder.

### Analysis

#### Membrane binding interface from CG simulations

The membrane-interacting residues of each protein were determined by computing the longest duration of interaction of each residue with the bilayer using the Prolint^76^ package.

The frequency of interaction for each amino acid and each secondary structure type in the LTD (Fig. 2d, 2e) were determined by summing the longest duration of interaction obtained from Prolint, for each amino acid/secondary structure type within the protein and obtaining normalized values between 0 and 1 for every protein. The mean interaction across all proteins was then determined by averaging over these normalized values.

#### Principal Component Analysis and Clustering from atomistic simulations

To analyze and compare the conformational changes that occur in the simulations of the apo and holo forms of each protein, we first performed Principal Component (PC) analysis of the protein’s Cα carbon atoms using the Scikit-learn package^77^ (version 0.21.3). The proteins were aligned, and the analysis was performed on the entire and concatenated set of simulations for each protein (6 apo, and 6 holo). The highly mobile N-and C-terminal ends of the proteins (2-5 residues, where necessary), as well as highly mobile residues 75 to 86 in STARD11 and residues 152 to 181 in Mdm12 were excluded from the analysis to avoid possible biasing of the PCs. After scrutiny of the resulting PCs, we focused our attention on the first PC i.e. PC1 (the PC with the highest variance), which showed most often a consistent convergence with the results obtained when the apo and holo simulations were analyzed separately. The projection of the protein dynamics along the analyzed PCs was further clustered using CLoNe^28^ and the relative apo and holo populations of each cluster were calculated by determining how many frames of each cluster were derived from the corresponding simulations.

#### Cavity volumes

Cavity volumes were estimated using the mdpocket tool of the fpocket package^78^ selecting only the internal protein cavity for the calculation.

Secondary structure images of the protein were rendered using VMD^79^ or ngl viewer^80^.

#### STARD11 experimental investigations Cell lines and culture conditions

HeLa cells were grown in DMEM high glucose, GlutaMAX™ (Gibco, USA) supplemented with 10% (v/v) foetal bovine serum (FBS), 4.5 g/L glucose, 2 mM L-glutamine, 1 U/mL penicillin and streptomycin under controlled atmosphere (5% CO_2_ and 95% air) at 37°C.

#### Plasmid transfection

Human STARD11 WT and mutants were inserted into pEGFP-C1 for WT or pcDNA3.1 eGFP vector for mutants to produce protein with eGFP at the N-terminus. Plasmids were transfected into HeLa cells with jetPRIME transfection reagent (Polyplus Transfection, 114-15) following the manufacturer’s instructions.

#### Antibodies

The following primary antibodies were used: rabbit anti-COL4A3BP (Sigma Aldrich, HPA035645, RRID: AB_10600700, 1:5,000 for WB, 1:300 for IF), rabbit anti-GOLPH3 (Abcam ab98023, RRID: AB_10860828, 1:300), mouse anti-GAPDH Clone 6C5 (Santa Cruz Biotechnology, sc-32233, RRID: AB_627679, 1:2,000). The following secondary antibodies were used: donkey A568-conjugated anti-rabbit (ThermoFisher Scientific, A-10042, RRID: AB_2534017, 1:400), donkey A647-conjugated anti-mouse (ThermoFisher Scientific, A-31571, RRID: AB_162542, 1:400 or Jackson ImmunoResearch, 715.605.150, AB_2340862, 1:200 for quantitative image analysis), donkey HRP-conjugated anti-rabbit (Jackson ImmunoResearch, 711-035-152, RRID: AB_10015282, 1:10,000), and donkey HRP-conjugated anti-mouse (Jackson ImmunoResearch, 715-035-150, RRID: AB_2340770). Hoechst was purchased from Life Technologies (H3570, 1:10,000 from 10mg/mL stock).

#### Immunofluorescence, staining and image analysis

HeLa cells were grown on glass coverslips, treated according to the experimental procedure, fixed with 4% paraformaldehyde for 15 min at RT and washed three times with PBS. After fixation, cells were blocked with 5% BSA and permeabilized with 0.5% saponin for 20 min at RT, followed by 1h incubation with selected antibodies against the antigen of interest in blocking solution. Cells were then washed three times with PBS and incubated with appropriate isotype-matched, AlexaFluor-conjugated secondary antibodies diluted in blocking solution for 30 min. After immunostaining, cells were washed three times in PBS and once in water, to remove salts. After Hoechst staining for nuclei, the samples were mounted with Fluoromount-G® (Southern Biotech, 0100-01) on glass microscope slides and analysed under a confocal microscope Leica SP8 with 63x oil objective (1.4 NA) or Zeiss LSM700 with 40x air objective (1.3 NA). Optical confocal sections were taken at 1 Airy unit under non-saturated conditions with a resolution of 1024×1024 pixels and frame average 4. Images were then processed using Fiji software (https://imagej.net/Fiji)^81^.

#### Quantitative cell imaging

HeLa cells were seeded in a μ-Plate 96 Well Black (IBIDI, 89626) at a concentration of 8 x 10^4^cell/well for 48h transfection experiment or 1.2×10^4^ cells/well for 16h transfection experiment. STARD11-GFP WT or mutant plasmids were transfected using the TransIT-X2® Transfection Reagent (Mirus Bio, MIR 6000). Cells were fixed in 3% paraformaldehyde for 20 min before staining with antibodies diluted in a PBS solution of 1% BSA and 0.05% saponin. After washing with an automated plate washer (BioTek EL406), cells were incubated for half an hour with appropriate secondary antibodies and nuclei were stained by Hoechst. Cells were left in PBS and imaged by ImageXpress® Micro Confocal microscope (Molecular Devices, Sunnyvale, CA); for each well, 49 frames were taken in widefield mode with a 40X objective. Images were quantified using MetaXpress Custom Module editor software to first segment the image and generate relevant masks, which were then applied on the fluorescent images to extract relevant measurements.

#### SDS-PAGE and Western blotting

After treatment, the cells were washed three times with PBS and lysed in a buffer consisting of 20 mM MOPS pH 7.0, 2 mM EGTA, 5 mM EDTA, 60 mM β-glycerophosphate, 30 mM NaF, 1 mM Na_3_VO_4_, 1% (v/v) Triton X-100, phosphatase inhibitor (PhosSTOP™, Sigma-Aldrich) and protease inhibitor (cOmplete™ Protease Inhibitor Cocktail, EDTA free, Roche). The lysates were clarified by centrifugation and quantified with Pierce™ BCA Protein Assay Kit (ThermoFisher) according to the manufacturer’s instructions. Samples were prepared by adding 4x SDS sample buffer, denatured at 95°C for 5 min and resolved by SDS-PAGE and immunoblot. For immunoblotting, the membrane strips containing the proteins of interest were blocked in TBS-T/ 5% BSA for 45 min at RT, and then incubated with the primary antibody diluted to its working concentration in the blocking buffer for 1h at RT. After washing with TBS-T, the strips were incubated for 1 h with the appropriate HRP-conjugated secondary antibody diluted in the blocking buffer. After washing with TBS-T, the strips were incubated with the ECL solution for 3 min and exposed to x-ray films, which were then scanned. The intensity of the bands and preparation of images was done using Fiji^81^ and Adobe Illustrator 2020.

Sphingolipidomics analysis carried out as described in ^82^. Frozen cell pellets were resuspended in 50 μL PBS and extracted with 1 mL Methanol/MTBE (methyl-tert-butyl ether)/Chloroform (MMC) [4:3:3; (v/v/v)] at 37°C (1400 rpm, 30 min). Internal lipid standards include D_7_SA (d18:0), D_7_SO (d18:1), dhCer (d18:0/12:0), ceramide (d18:1/12:0), glucosylceramide (d18:1/8:0), SM (d18:1/18:1(D_9_)), and D_7_-S1P. The single-phase supernatant was collected, dried under N_2_ and dissolved in 70 μL methanol. Untargeted lipid analysis was performed on a high-resolution Q Exactive MS analyzer (Thermo Scientific) as described earlier^83^.

### Mdm12 experimental investigations

#### Yeast studies

Strains, plasmids and primers used in this study are listed in Supplementary Table 1. Yeast cells were cultured at 30 °C in synthetic defined (SD) medium with 2% glucose (2% glucose, 0.5% NH_4_-sulfate, 0.17% yeast nitrogen base and amino acids). The MDM12 gene corresponding to the L28C L244C mutant was synthesized at GenScript, Netherlands.

#### Microscopy

Cells were grown in SD-leucine-uracil for the selection of the Mdm12-GFP plasmid and the mitochondria matrix-targeted CFAse mcherry plasmid. Images were acquired using a Nikon Widefield microscope equipped with a 100x NA 1.45 objective lens, a Spectra III Light Engine illumination and an ORCA Fusion BT camera. Image acquisition was done at room temperature. Images were processed further using the FIJI ImageJ bundle (version 1.53c). For quantification of mitochondria morphology, filenames were anonymized before determining the number of cells with the indicated morphology, using the Cell Counter plug-in in FIJI.

#### Pulse-labeling, lipid extraction and MS analysis

The following steps were carried out as previously described^43^. Pre-cultures in SD medium were diluted to 0.7 OD_600_ ml^−1^ in 25 ml and treated with 0.5 mM auxin for 7 h. Next, cells were pulse-labelled with 2 mM d-methionine and grown at 30 °C. At the indicated timepoints, 8 OD_600_ of cells was pelleted, snap-frozen and stored at −80 °C. Lipids were extracted as described previously with minor modifications^84^. Briefly, cells were washed in ice-cold water and subsequently resuspended in 1.5 ml of extraction solvent containing ethanol, water, diethyl ether, pyridine and 4.2 N ammonium hydroxide (v/v 15:15:5:1:0.18). After the addition of 300 µl glass beads, samples were vortexed vigorously for 5 min and incubated at 60 °C for 20 min. Cell debris were pelleted by centrifugation at 1,800*g* for 10 min, and the supernatant was dried under a stream of nitrogen. The dried extract was resuspended in 1 ml of water-saturated butanol and sonicated for 5 min in a water bath sonicator. Then, 500 µl of water was added and vortexed further for 2 min. After centrifugation at 3,000*g*, the upper butanol phase was collected, dried under a stream of nitrogen and resuspended in 50% methanol for lipidomics analysis.

LC analysis was performed as described previously with several modifications^85^. Phospholipids were separated on a nanoAcquity ultra-performance liquid chromatography unit (Waters) equipped with a HSS T3 capillary column (150 m × 30 mm, 1.8 m particle size; Waters), applying a 10 min linear gradient of buffer A (5 mM ammonium acetate in acetonitrile:water 60:40) and B (5 mM ammonium acetate in isopropanol:acetonitrile 90:10) from 10% B to 100% B. Conditions were kept at 100% B for the next 7 min, followed by a 8 min re-equilibration to 10% B. The injection volume was 1 µl. The flow rate was constant at 2.5 µl min^−1^.

The ultra-performance liquid chromatography unit was coupled to QExactive mass spectrometer (Thermo) by a nanoESI source (New Objective Digital PicoView 550) equipped with the Thermo QExactive XCalibur software (version 4.0.27.10). The source was operated with a spray voltage of 2.9 kV in positive mode and 2.5 kV in negative mode. Sheath gas flow rate was set to 25 and 20 for positive and negative mode, respectively. MS data were acquired using either positive or negative polarization, alternating between full MS and all-ion-fragmentation scans. Full scan MS spectra were acquired in profile mode from 107 *m*/*z* to 1,600 *m*/*z* with an automatic gain control target of 1 × 10^6^, an Orbitrap resolution of 70,000 and a maximum injection time of 200 ms. All-ion-fragmentation spectra were acquired from 107 *m*/*z* to 1,600 *m*/*z* with an automatic gain control value of 5 × 10^4^, a resolution of 17,500 and a maximum injection time of 50 ms, and fragmented with a normalized collision energy of 20, 30 and 40 (arbitrary units). Generated fragment ions were scanned in the linear trap. Positive ion mode was employed for monitoring PC and negative ion mode was used for monitoring PS and PE. Lipid species were identified on the basis of their *m*/*z* and elution time. We used a standard mixture comprising PS 10:0/10:0, PE 17:0/17:0, PC 17:0/17:0, PG 17:0/17:0 and PI 12:0/13:0 for deriving an estimate of specific elution times. Lipid intensities were quantified using the Skyline (version 21.2.0.369) software^86^. For each phospholipid, signal was integrated for the precursor species (m), cyclopropane species (m_+14_) and species that appear upon pulse labelling with d-methionine (m_+9_, m_+16_, m_+23_ and m_+25_). Fraction of labelled headgroups was calculated as (m_+9_ + m_+23_ + m_+25_) / (m + m_+9_ + m_+14_ + m_+16_ + m_+23_ + m_+25_).

Fraction of labelled cyclopropane was calculated as (m_+16_ + m_+25_)/(m + m_+9_ + m_+14_ + m_+16_ + m_+23_ + m_+25_). Fraction of labeled headgroups and cyclopropane, independent of transport was calculated as (m_+9_ + m_+23_)/(m + m_+9_ + m_+14_ + m_+16_ + m_+23_ + m_+25_) and (m_+16_)/(m_+14_ + m_+16_ + m_+23_ + m_+25_), respectively. Fraction of doubly labelled mass-tagged species was calculated as (m_+25_)/(m + m_+9_ + m_+14_ + m_+16_ + m_+23_ + m_+25_).

## Data availability

All input files for atomistic and coarse-grained MD simulations, structures of the LTD with the membrane binding interface, and representative apo-like and holo-like conformations of each LTD arising from clustering of the atomistic simulations can be found at: https://doi.org/10.5281/zenodo.7819506

## Acknowledgments

S.V. acknowledges support by the SNSF (PP00P3_194807) and by the European Research Council under the European Union’s Horizon 2020 research and innovation program (grant agreement no. 803952). G.D’A. acknowledges support by the Swiss Cancer League, KFS-4999-02-2020; by the EPFL institutional fund; and by SNSF (310030_184926); This work was supported by grants from the Swiss National Supercomputing Centre under projects ID s1030 and s1132. S.V. and A.J.P. acknowledge support from the Novartis Forschungsstiftung via a FreeNovation grant. M.A.L. acknowledges support from the Foundation Suisse de Recherche sur le Maladies Musculaires (FSRMM). Lipidomics measurements for Mdm12 were performed at the Functional Genomics Center Zurich (FGCZ). We especially thank Sebastian Streb of the FGCZ Metabolomics division for excellent technical guidance. The authors thank P. Campomanes for critical reading of the manuscript. We thank Dimitri Moreau and Stefania Vossio (ACCESS Geneva) for microscopy and data analysis.

## Supplementary Information

**Supplementary Table 1.** Plasmids used in this study.

| Plasmid | Genotype | Reference |
| --- | --- | --- |
| pAJ15 | pRS416-TEFpr-Su9(ss)-CFase-mCHERRY | Ref <sup>43</sup> |
| pAJ16 | pRS415-Mdm12pr-Mdm12 | Ref <sup>43</sup> |
| pAJ17 | pRS415-Mdm12pr-Mdm12 (L28C L244C) | Ref <sup>43</sup> |
| pAJ18 | pRS415-Mdm12pr-Mdm12-GFP | Ref <sup>43</sup> |
| pAJ19 | pRS415-Mdm12pr-Mdm12 (L28C L244C)-GFP | Ref <sup>43</sup> |

**Figure S1.**
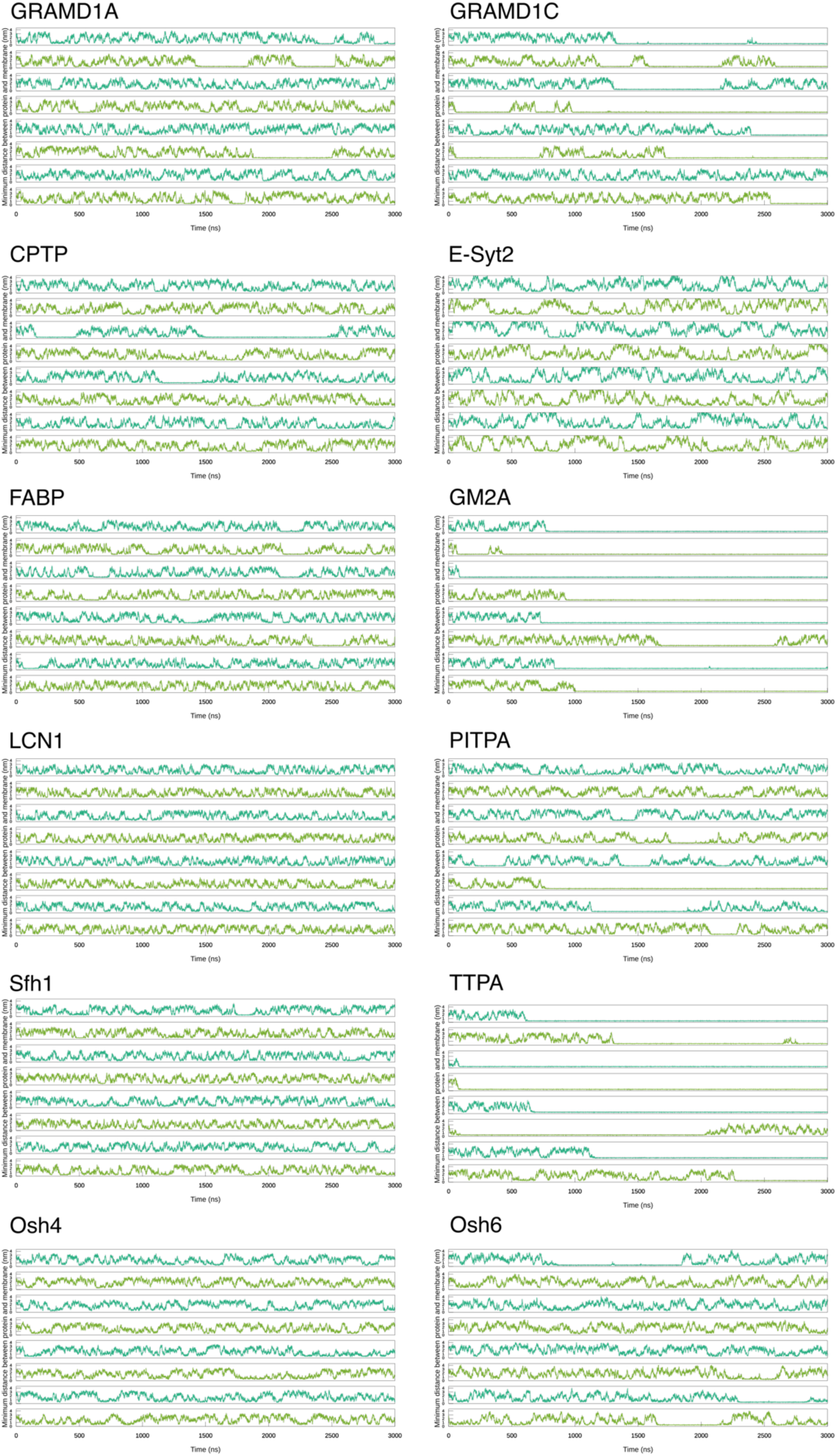
Time trace of minimum distance values between the protein and the bilayer, for each replica of simulation; indicates transient and reversible interactions.

**Figure S2.**
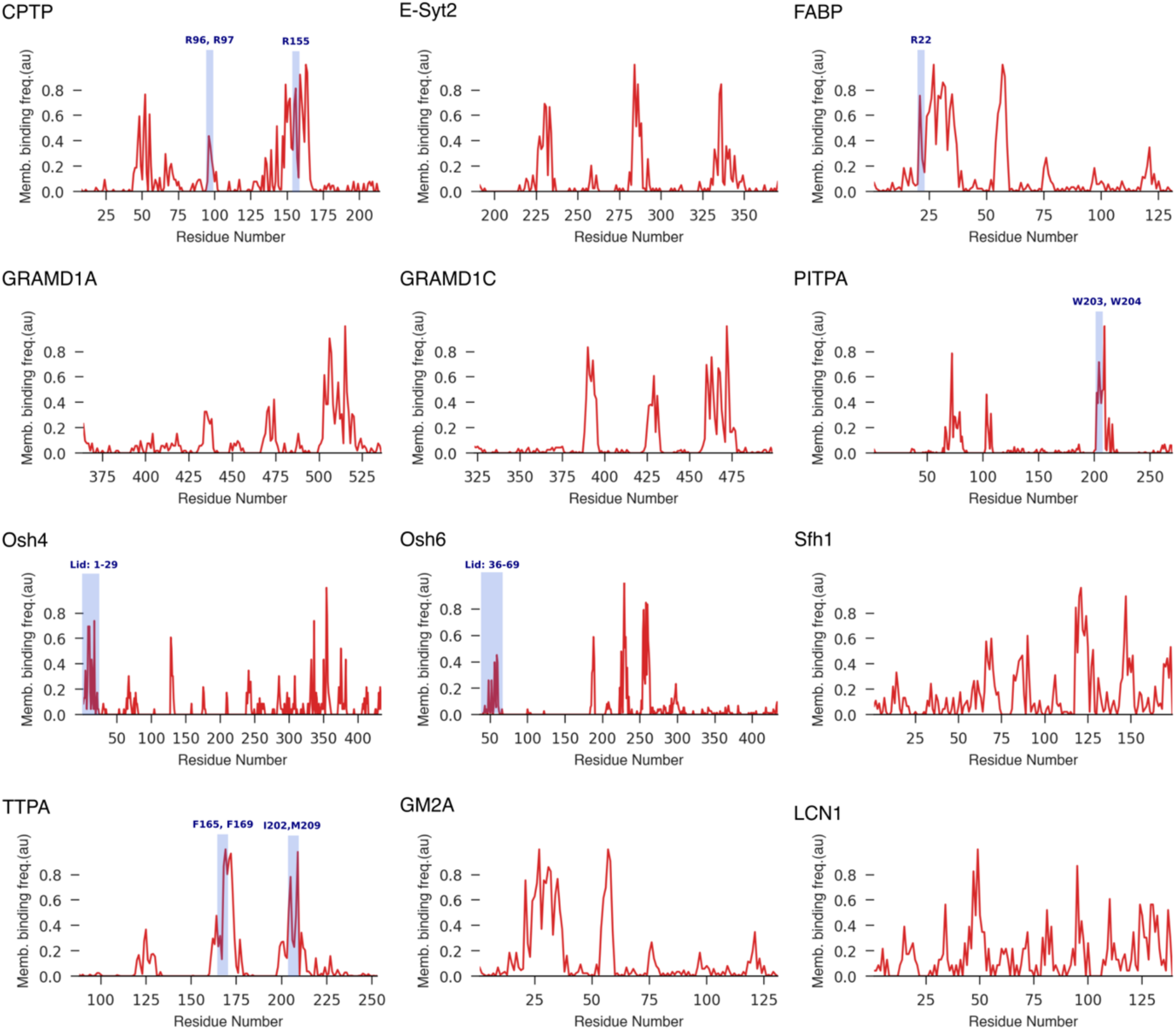
Residue-wise frequency of interaction with the lipid bilayer. Residues that have been experimentally proposed to be crucial for membrane binding are highlighted in blue.

**Figure S3.**
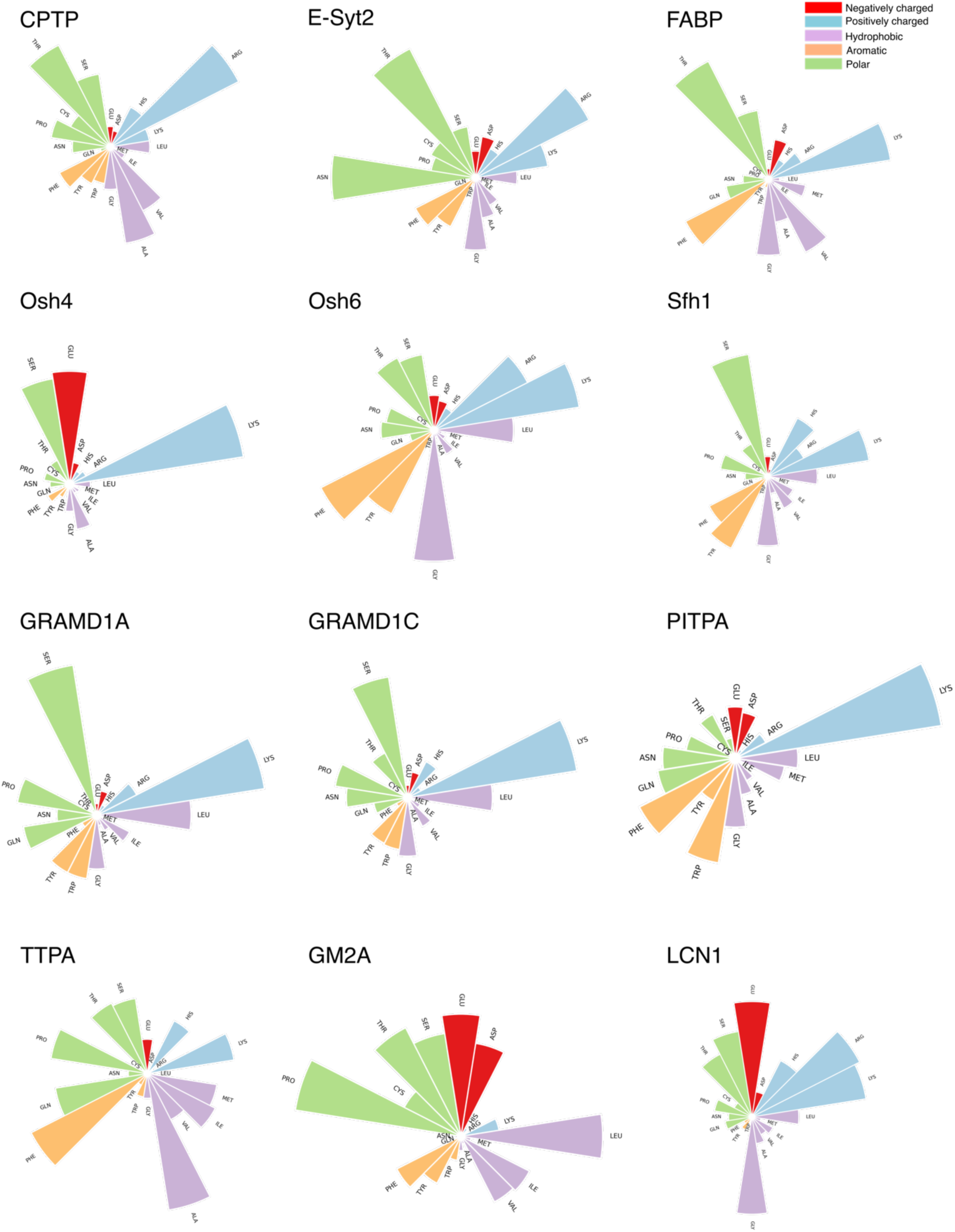
Interaction frequency of each amino acid with the lipid bilayer, shown for each LTP in our dataset.

**Figure S4.**
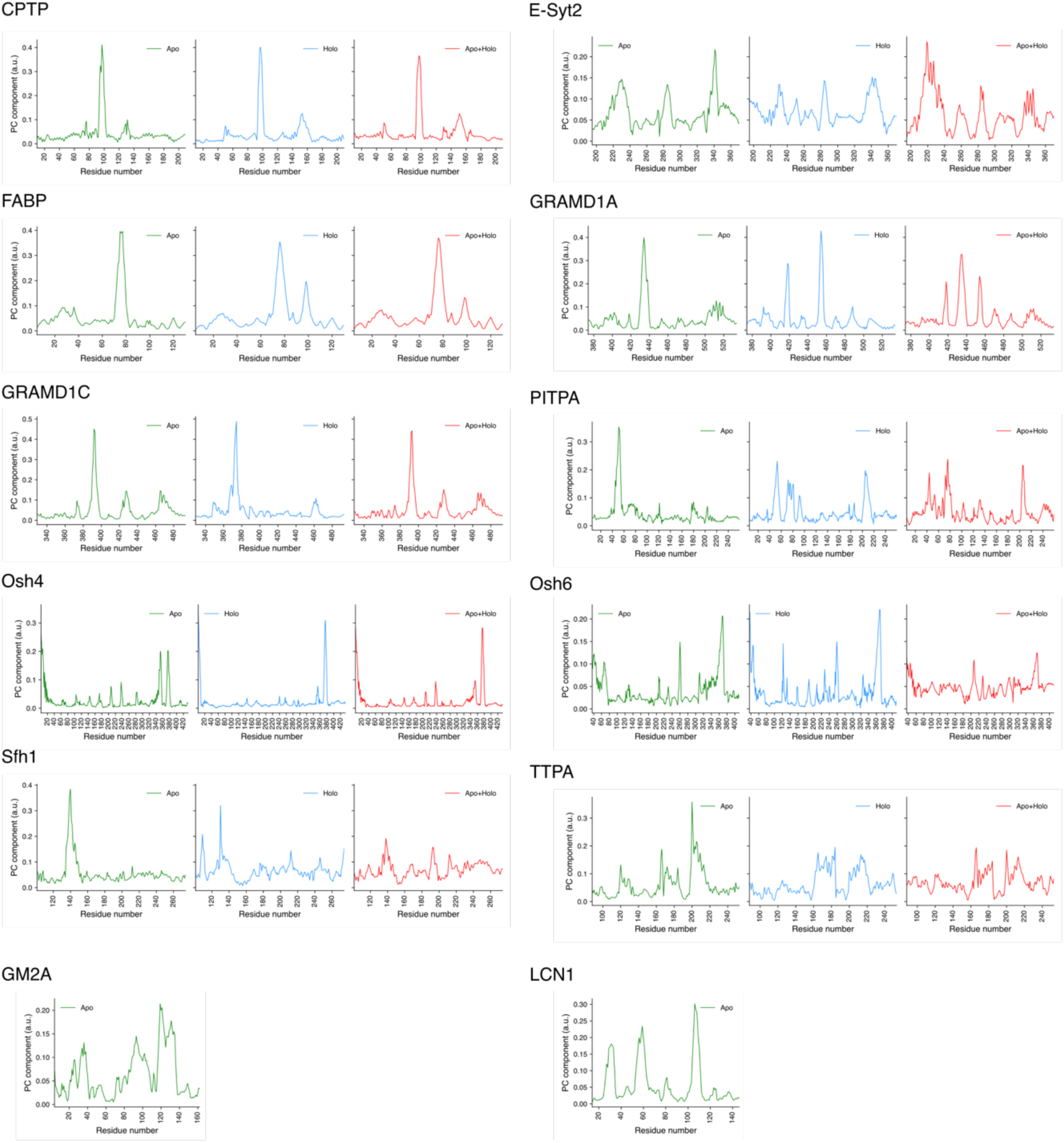
Residue-wise contribution to principal component PC1 for the apo (green), holo (blue), and combined (red) trajectories. GM2A and LCN1 were simulated in the apo-form alone due to the lack of a lipid-bound crystal structure of the protein.

**Figure S5.**
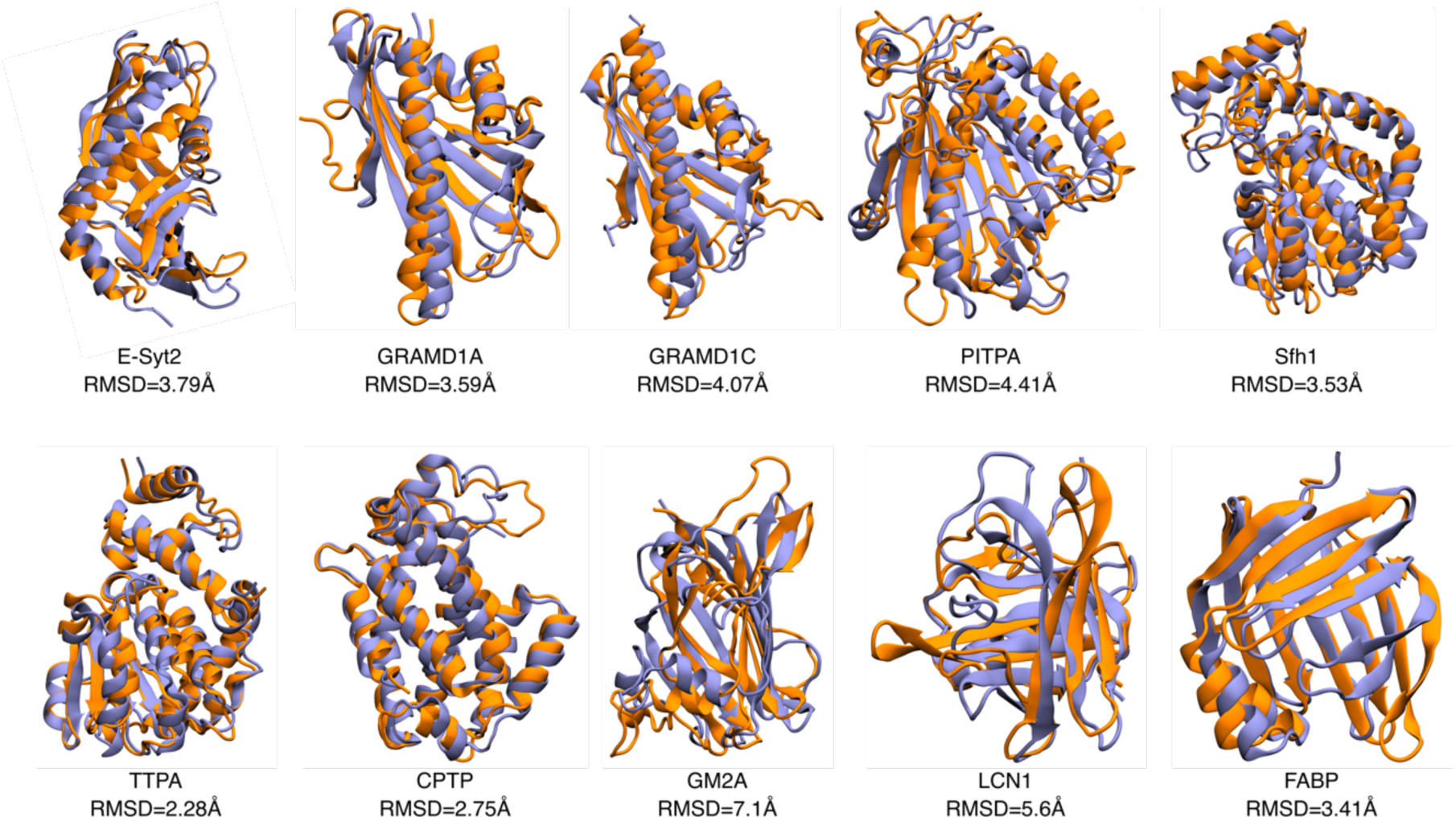
Comparison of the apo-like (orange) and holo-like (purple) structures of the LTDs arising from the extreme ends of the clustering procedure, and the RMSD between them. Osh4 and Osh6 not shown as the N-terminal lid of the protein that exhibits the largest motion can exist in several folded and unfolded states resulting in a wide range of RMSD values.

**Figure S6.**
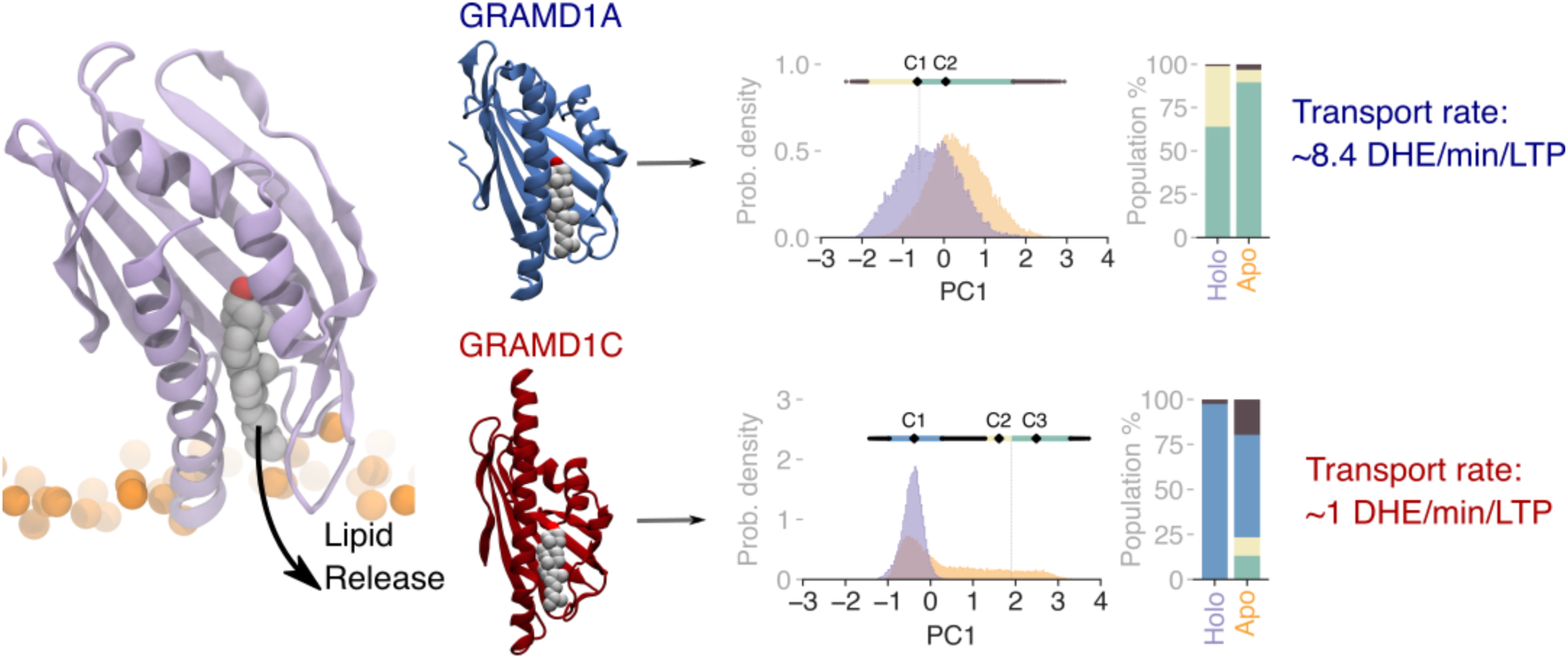
Despite their structural similarities, the difference in the dynamical behaviour of GRAMD1A and GRAMD1C could explain their difference in transport rates.

**Figure S7.**
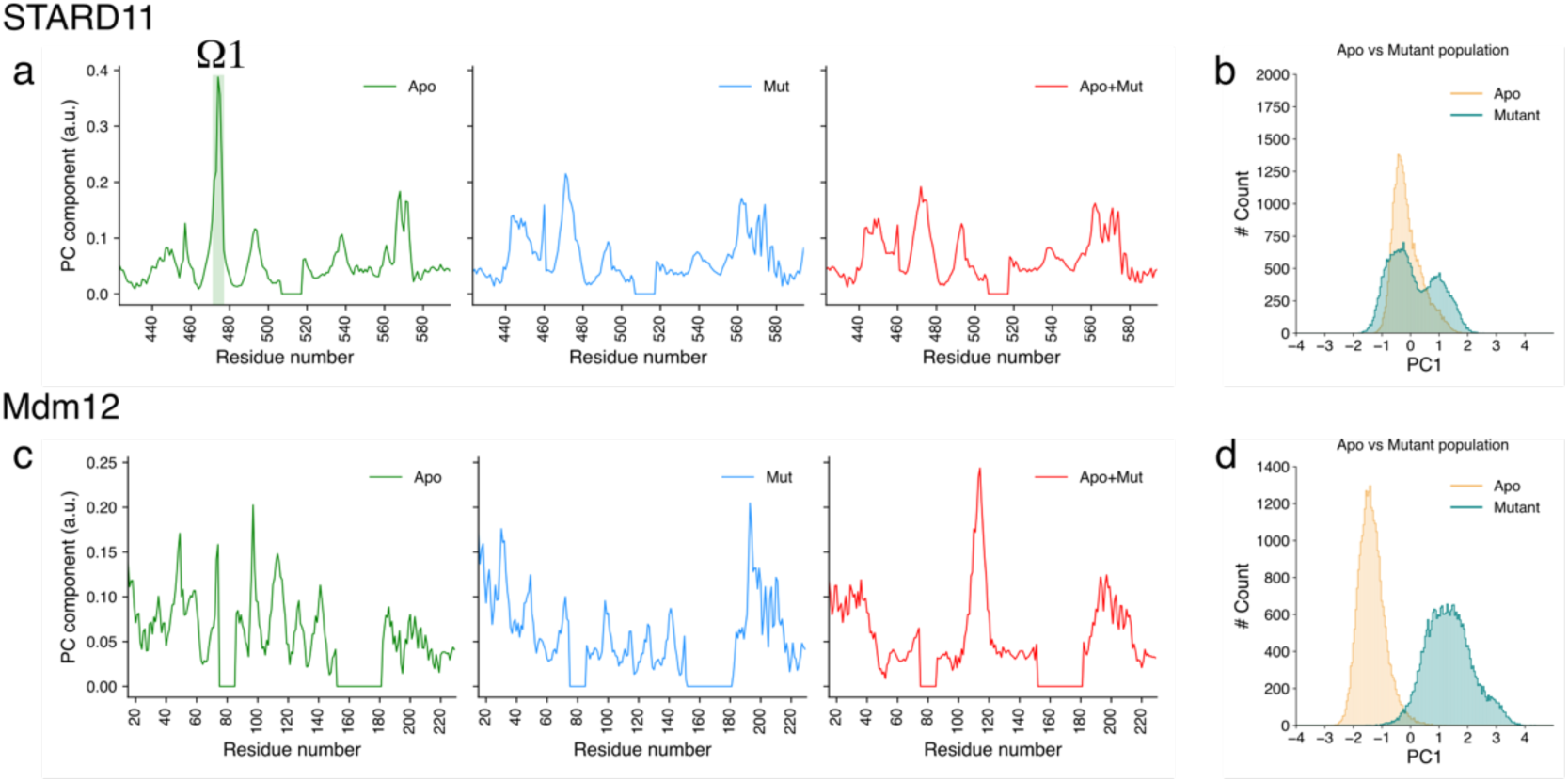
PCA of STARD11 and Mdm12 + mutants. Residue-wise contribution to principal component PC1 for the apo (green), mutant (blue), and combined (red) trajectories for STARD11 (a) and Mdm12 (c) and their population distributions of PC1 from simulations of apo form (orange), mutant form (blue) for STARD11 (b) and Mdm12 (d).

**Figure S8.**
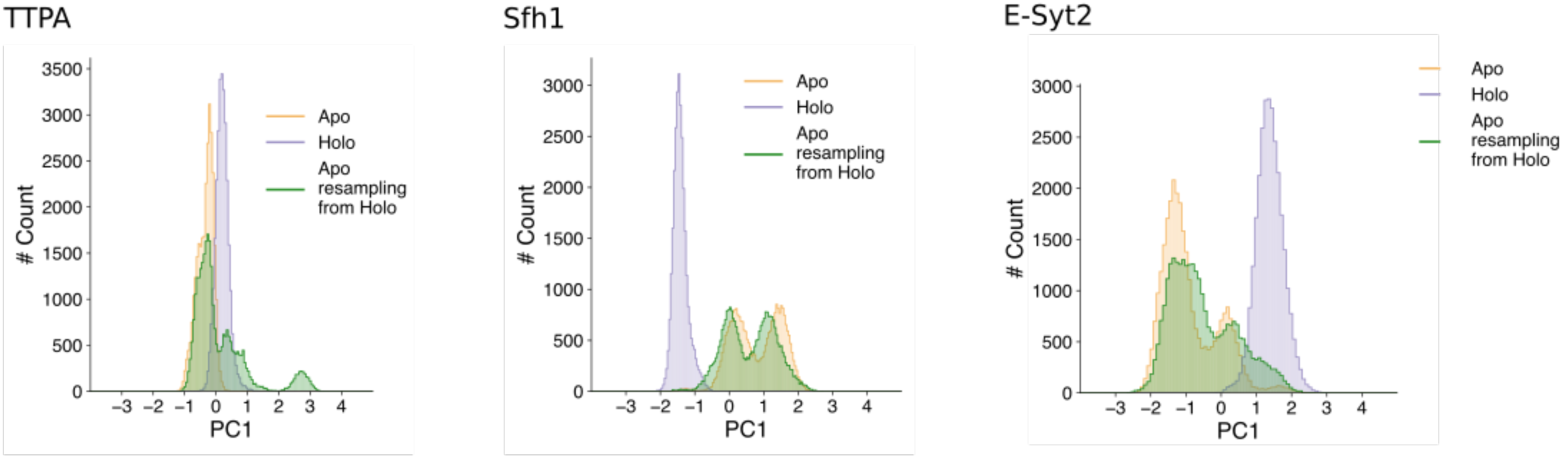
The population distributions of PC1 from simulations of apo form (orange), holo form (purple), and simulations of the protein in the lipidless-form starting from the final structure obtained from the holo simulations by removing the lipid from the last frame of the simulations (green). The PC obtained from resampling (green) are similar to the ones obtained from simulating the apo form of the LTD starting from the crystal structure (orange), indicating that all conformational changes observed are reversible and the protein is not trapped in a metastable state.

## References

1. G. van Meer, D. R. Voelker and G.W. Feigenson, Nature Reviews Molecular Cell Biology, 2008, 9, 112–124.

2. H. Sunshine and M. L. Iruela-Arispe, Curr Opin Lipidol, 2017, 28, 408–413.

3. P. Fagone and S. Jackowski, J Lipid Res, 2009, 50 Suppl, S311–316.

4. J. S. Bonifacino and B. S. Glick, Cell, 2004, 116, 153–166.

5. K. M. Reinisch and W.A. Prinz, Journal of Cell Biology, 2021, 220.

6. L. H. Wong, A. Copic and T.P. Levine, Trends in biochemical sciences, 2017, 42, 516–530.

7. A. Chiapparino, K. Maeda, D. Turei, J. Saez-Rodriguez and A. C. Gavin, Prog Lipid Res, 2016, 61, 30–39.

8. A.-E. Saliba, I. Vonkova, S. Deghou, S. Ceschia, C. Tischer, K. G. Kugler, P. Bork, J. Ellenberg and A.-C. Gavin, Nature Protocols, 2016, 11, 1021–1038.

9. Louise H. Wong and Tim P. Levine, Biochemical Society Transactions, 2016, 44, 517–527.

10. N. F. Lipp, R. Gautier, M. Magdeleine, M. Renard, V. Albanèse, A. Čopič and G. Drin, Nat Commun, 2019, 10, 3926.

11. D. K. Simanshu, R. K. Kamlekar, D. S. Wijesinghe, X. Zou, X. Zhai, S. K. Mishra, J. G. Molotkovsky, L. Malinina, E. H. Hinchcliffe, C. E. Chalfant, R. E. Brown and D. J. Patel, Nature, 2013, 500, 463–467.

12. X. Miliara, T. Tatsuta, J.-L. Berry, S. L. Rouse, K. Solak, D. S. Chorev, D. Wu, C. V. Robinson, S. Matthews and T. Langer, Nature Communications, 2019, 10, 1130.

13. J. Moser von Filseck, A. Čopič, V. Delfosse, S. Vanni, C. L. Jackson, W. Bourguet and G. Drin, Science, 2015, 349, 432–436.

14. J. Chung, F. Torta, K. Masai, L. Lucast, H. Czapla, L. B. Tanner, P. Narayanaswamy, M. R. Wenk, F. Nakatsu and P. De Camilli, Science, 2015, 349, 428–432.

15. S. Srinivasan, V. Zoni and S. Vanni, Faraday Discussions, 2021, DOI: 10.1039/D0FD00058B.

16. Y.-G. Gao, X. Zhai, I. A. Boldyrev, J. G. Molotkovsky, D. J. Patel, L. Malinina and R.E. Brown, Journal of Biological Chemistry, 2021, 296.

17. F. M. Herr, J. Aronson and J. Storch, Biochemistry, 1996, 35, 1296–1303.

18. M. de Saint-Jean, V. Delfosse, D. Douguet, G. Chicanne, B. Payrastre, W. Bourguet, B. Antonny and G. Drin, J Cell Biol, 2011, 195, 965–978.

19. S. Shadan, R. Holic, N. Carvou, P. Ee, M. Li, J. Murray-Rust and S. Cockcroft, Traffic, 2008, 9, 1743–1756.

20. W. X. Zhang, V. Thakur, A. Lomize, I. Pogozheva, C. Panagabko, M. Cecchini, M. Baptist, S. Morley, D. Manor and J. Atkinson, Journal of molecular biology, 2011, 405, 972–988.

21. A. Mulgrew-Nesbitt, K. Diraviyam, J. Wang, S. Singh, P. Murray, Z. Li, L. Rogers, N. Mirkovic and D. Murray, Biochimica et biophysica acta, 2006, 1761, 812–826.

22. M. A. Lemmon and K. M. Ferguson, Biochem J, 2000, 350 Pt 1, 1–18.

23. Hanif M. Khan, T. He, E. Fuglebakk, C. Grauffel, B. Yang, Mary F. Roberts, A. Gershenson and N. Reuter, Biophysical Journal, 2016, 110, 1367–1378.

24. J. E. Johnson and R. B. Cornell, Molecular Membrane Biology, 1999, 16, 217–235.

25. R. V. Stahelin, F. Long, B. J. Peter, D. Murray, P. De Camilli, H. T. McMahon and W. Cho, Journal of Biological Chemistry, 2003, 278, 28993–28999.

26. S. Hayward and N. Go, Annual Review of Physical Chemistry, 1995, 46, 223–250.

27. M. A. Balsera, W. Wriggers, Y. Oono and K. Schulten, The Journal of Physical Chemistry, 1996, 100, 2567–2572.

28. S. Träger, G. Tamò, D. Aydin, G. Fonti, M. Audagnotto and M. Dal Peraro,Bioinformatics, 2020, DOI: 10.1093/bioinformatics/btaa742.

29. S. Kumar, B. Ma, C. J. Tsai, N. Sinha and R. Nussinov, Protein Sci, 2000, 9, 10–19.

30. D. D. Boehr, R. Nussinov and P. E. Wright, Nature Chemical Biology, 2009, 5, 789–796.

31. F. A. Horenkamp, D. P. Valverde, J. Nunnari and K. M. Reinisch, The EMBO Journal, 2018, 37, e98002.

32. Y. Tsujishita and J. H. Hurley, Nat Struct Biol, 2000, 7, 408–414.

33. M. R. Wozny, A. Di Luca, D. R. Morado, A. Picco, P. C. Hoffmann, E. A. Miller, S. Vanni and W. Kukulski, bioRxiv, 2022, DOI: 10.1101/2022.04.12.488000,2022.2004.2012.488000.

34. B. Kornmann, E. Currie, S. R. Collins, M. Schuldiner, J. Nunnari, J. S. Weissman and P. Walter, Science, 2009, 325, 477–481.

35. M. Kawano, K. Kumagai, M. Nishijima and K. Hanada, The Journal of biological chemistry, 2006, 281, 30279–30288.

36. K. Hanada, K. Kumagai, S. Yasuda, Y. Miura, M. Kawano, M. Fukasawa and M. Nishijima, Nature, 2003, 426, 803–809.

37. K. Hanada, Proc Jpn Acad Ser B Phys Biol Sci, 2010, 86, 426–437.

38. T. Naito, B. Ercan, L. Krshnan, A. Triebl, D. H. Z. Koh, F.-Y. Wei, K. Tomizawa, F. T. Torta, M. R. Wenk and Y. Saheki, eLife, 2019, 8, e51401.

39. C. Gehin, M. A. Lone, W. Lee, L. Capolupo, S. Ho, A. M. Adeyemi, E. H. Gerkes, A P. Stegmann, E. López-Martín, E. Bermejo-Sánchez, B. Martínez-Delgado, C. Zweier, C. Kraus, B. Popp, V. Strehlow, D. Gräfe, I. Knerr, E. R. Jones, S. Zamuner, L. A. Abriata, V. Kunnathully, B. E. Moeller, A. Vocat, S. Rommelaere, J. P. Bocquete, E. Ruchti, G. Limoni, M. Van Campenhoudt, S. Bourgeat, P. Henklein, C. Gilissen, B. W. van Bon, R. Pfundt, M. H. Willemsen, J. H. Schieving, E. Leonardi, F. Soli, A. Murgia, H. Guo, Q. Zhang, K. Xia, C. R. Fagerberg, C. P. Beier, M. J. Larsen, I. Valenzuela, P. Fernández-Álvarez, S. Xiong, R. Śmigiel, V. López-González, L. Armengol, M. Morleo, A. Selicorni, A. Torella, M. Blyth, N. S. Cooper, V. Wilson, R. Oegema, Y. Herenger, A. Garde, A. L. Bruel, F. Tran Mau-Them, A. B. Maddocks, J. M. Bain, M. A. Bhat, G. Costain, P. Kannu, A. Marwaha, N. L. Champaigne, M. J. Friez, E. B. Richardson, V. K. Gowda, V. M. Srinivasan, Y. Gupta, T. Y. Lim, S. Sanna-Cherchi, B. Lemaitre, T. Yamaji, K. Hanada, J. E. Burke, A. M. Jakšić, B. D. McCabe, P. De Los Rios, T. Hornemann, G. D’Angelo and V. A. Gennarino, J Clin Invest, 2023, DOI: 10.1172/jci165019.

40. K. Kumagai, M. Kawano, F. Shinkai-Ouchi, M. Nishijima and K. Hanada, The Journal of biological chemistry, 2007, 282, 17758–17766.

41. T. Fugmann, A. Hausser, P. Schöffler, S. Schmid, K. Pfizenmaier and M. A. Olayioye, J Cell Biol, 2007, 178, 15–22.

42. S. Capasso, L. Sticco, R. Rizzo, M. Pirozzi, D. Russo, N. A. Dathan, F. Campelo, J. van Galen, M. Hölttä-Vuori, G. Turacchio, A. Hausser, V. Malhotra, I. Riezman, H. Riezman, E. Ikonen, C. Luberto, S. Parashuraman, A. Luini and G. D’Angelo, Embo j, 2017, 36, 1736–1754.

43. A. T. John Peter, C. Petrungaro, M. Peter and B. Kornmann, Nature Cell Biology, 2022, 24, 996–1004.

44. K. S. Dimmer, S. Fritz, F. Fuchs, M. Messerschmitt, N. Weinbach, W. Neupert and B. Westermann, Mol Biol Cell, 2002, 13, 847–853.

45. Y. G. Gao, X. Zhai, I. A. Boldyrev, J. G. Molotkovsky, D. J. Patel, L. Malinina and R. E. Brown, The Journal of biological chemistry, 2021, 296, 100600.

46. W. X. Zhang, V. Thakur, A. Lomize, I. Pogozheva, C. Panagabko, M. Cecchini, M. Baptist, S. Morley, D. Manor and J. Atkinson, Journal of molecular biology, 2011, 405, 972–988.

47. H. Saaren-Seppälä, M. Jauhiainen, T. M. Tervo, B. Redl, P. K. Kinnunen and J. M. Holopainen, Invest Ophthalmol Vis Sci, 2005, 46, 3649–3656.

48. B. Rogaski, J. B. Lim and J.B. Klauda, The Journal of Physical Chemistry B, 2010, 114, 13562–13573.

49. G. Khelashvili, N. Chauhan, K. Pandey, D. Eliezer and A. K. Menon, eLife, 2019, 8, e53444.

50. L. H. Wong, A. T. Gatta and T.P. Levine, Nature Reviews Molecular Cell Biology, 2019, 20, 85–101.

51. J. C. Holthuis and T.P. Levine, Nat Rev Mol Cell Biol, 2005, 6, 209–220.

52. J. S. Dittman and A. K. Menon, Trends in biochemical sciences, 2017, 42, 90–97.

53. J. Moser von Filseck, A. Čopič, V. Delfosse, S. Vanni, C. L. Jackson, W. Bourguet and G. Drin, Science, 2015, 349, 432.

54. J. M. von Filseck, S. Vanni, B. Mesmin, B. Antonny and G. Drin, Nature Communications, 2015, 6, 6671.

55. A. Grabon, A. Orłowski, A. Tripathi, J. Vuorio, M. Javanainen, T. Róg, M. Lönnfors, M. I. McDermott, G. Siebert, P. Somerharju, I. Vattulainen and V. A. Bankaitis, The Journal of biological chemistry, 2017, 292, 14438–14455.

56. Y. M. Huang, Y. Miao, J. Munguia, L. Lin, V. Nizet and J. A. McCammon, Protein Sci, 2016, 25, 1430–1437.

57. X. Zhang, H. Xie, D. Iaea, G. Khelashvili, H. Weinstein and F. R. Maxfield, Journal of Biological Chemistry, 2022, 298.

58. N. Kudo, K. Kumagai, R. Matsubara, S. Kobayashi, K. Hanada, S. Wakatsuki and R. Kato, Journal of molecular biology, 2010, 396, 245–251.

59. J. Dong, X. Du, H. Wang, J. Wang, C. Lu, X. Chen, Z. Zhu, Z. Luo, L. Yu, A. J. Brown, H. Yang and J.-W. Wu, Nature Communications, 2019, 10, 829.

60. H. Wang, Q. Ma, Y. Qi, J. Dong, X. Du, J. Rae, J. Wang, W.-F. Wu, A. J. Brown, R. G. Parton, J.-W. Wu and H. Yang, Molecular Cell, 2019, 73, 458–473.e457.

61. J. G. Alb, A. Gedvilaite, R. T. Cartee, H. B. Skinner and V.A. Bankaitis, Proc Natl Acad Sci U S A, 1995, 92, 8826–8830.

62. X. Periole, M. Cavalli, S.-J. Marrink and M. A. Ceruso, Journal of Chemical Theory and Computation, 2009, 5, 2531–2543.

63. H. M. Berman, J. Westbrook, Z. Feng, G. Gilliland, T. N. Bhat, H. Weissig, I. N. Shindyalov and P. E. Bourne, Nucleic acids research, 2000, 28, 235–242.

64. Y. Qi, H. I. Ingólfsson, X. Cheng, J. Lee, S. J. Marrink and W. Im, Journal of Chemical Theory and Computation, 2015, 11, 4486–4494.

65. D. Van Der Spoel, E. Lindahl, B. Hess, G. Groenhof, A. E. Mark and H. J. C. Berendsen, Journal of computational chemistry, 2005, 26, 1701–1718.

66. P. C. T. Souza, R. Alessandri, J. Barnoud, S. Thallmair, I. Faustino, F. Grünewald, I. Patmanidis, H. Abdizadeh, B. M. H. Bruininks, T. A. Wassenaar, P. C. Kroon, J. Melcr, V. Nieto, V. Corradi, H. M. Khan, J. Domański, M. Javanainen, H. Martinez-Seara, N. Reuter, R. B. Best, I. Vattulainen, L. Monticelli, X. Periole, D. P. Tieleman, A. H. de Vries and S. J. Marrink, Nature Methods, 2021, 18, 382–388.

67. G. Bussi, D. Donadio and M. Parrinello, The Journal of Chemical Physics, 2007, 126, 014101.

68. M. Parrinello and A. Rahman, Journal of Applied Physics, 1981, 52, 7182–7190.

69. I. G. Tironi, R. Sperb, P. E. Smith and W. F. van Gunsteren, The Journal of chemical physics, 1995, 102, 5451–5459.

70. D. H. de Jong, S. Baoukina, H. I. Ingólfsson and S. J. Marrink, Computer Physics Communications, 2016, 199, 1–7.

71. H. Jeong, J. Park and C. Lee, EMBO reports, 2016, 17, 1857–1871.

72. Y. Qi, X. Cheng, W. Han, S. Jo, K. Schulten and W. Im, Journal of Chemical Information and Modeling, 2014, 54, 1003–1009.

73. C. M. Schauder, X. Wu, Y. Saheki, P. Narayanaswamy, F. Torta, M. R. Wenk, P. De Camilli and K. M. Reinisch, Nature, 2014, 510, 552–555.

74. K. Vanommeslaeghe, E. P. Raman and A. D. MacKerell, Jr., J Chem Inf Model, 2012, 52, 3155–3168.

75. J. Huang, S. Rauscher, G. Nawrocki, T. Ran, M. Feig, B. L. de Groot, H. Grubmüller and A. D. MacKerell, Jr., Nature methods, 2017, 14, 71–73.

76. B. I. Sejdiu and D. P. Tieleman, Nucleic acids research, 2021, 49, W544–W550.

77. F. Pedregosa, G. Varoquaux, A. Gramfort, V. Michel, B. Thirion, O. Grisel, M. Blondel, P. Prettenhofer, R. Weiss and V. Dubourg, the Journal of machine Learning research, 2011, 12, 2825–2830.

78. P. Schmidtke, A. Bidon-Chanal, F. J. Luque and X. Barril, Bioinformatics, 2011, 27, 3276–3285.

79. W. Humphrey, A. Dalke and K. Schulten, Journal of Molecular Graphics, 1996, 14, 33–38.

80. A. S. Rose and P. W. Hildebrand, Nucleic acids research, 2015, 43, W576–W579.

81. J. Schindelin, I. Arganda-Carreras, E. Frise, V. Kaynig, M. Longair, T. Pietzsch, S. Preibisch, C. Rueden, S. Saalfeld, B. Schmid, J. Y. Tinevez, D. J. White, V. Hartenstein, K. Eliceiri, P. Tomancak and A. Cardona, Nat Methods, 2012, 9, 676–682.

82. M. A. Lone, A. J. Hülsmeier, E. M. Saied, G. Karsai, C. Arenz, A. von Eckardstein and T. Hornemann, Proc. Natl. Acad. Sci. U. S. A., 2020, 117, 15591–15598.

83. M. A. Lone, M. J. Aaltonen, A. Zidell, H. F. Pedro, J. A. Morales Saute, S. Mathew, P. Mohassel, C. G. Bönnemann, E. A. Shoubridge and T. Hornemann, The Journal of Clinical Investigation, 2022, 132.

84. A. X. da Silveira Dos Santos, I. Riezman, M. A. Aguilera-Romero, F. David, M. Piccolis, R. Loewith, O. Schaad and H. Riezman, Mol Biol Cell, 2014, 25, 3234–3246.

85. J. M. Castro-Perez, J. Kamphorst, J. DeGroot, F. Lafeber, J. Goshawk, K. Yu, J. P. Shockcor, R. J. Vreeken and T. Hankemeier, Journal of Proteome Research, 2010, 9, 2377–2389.

86. K. J. Adams, B. Pratt, N. Bose, L. G. Dubois, L. St John-Williams, K. M. Perrott, K. Ky, P. Kapahi, V. Sharma, M. J. MacCoss, M. A. Moseley, C. A. Colton, B. X. MacLean, B. Schilling and J. W. Thompson, J Proteome Res, 2020, 19, 1447–1458.

